# Cell type-resolved chromatin accessibility clocks for brain aging

**DOI:** 10.64898/2026.08.22.746439

**Authors:** Patrick Z. Yu, Doudou Yu, Yao Xue, William Stafford Noble

## Abstract

Aging is a progressive decline in biological function that is proposed to be driven by the accumulation of epigenetic noise and the loss of epigenetic information. Among epigenetic readouts, DNA methylation has been extensively used to develop aging clocks, machine learning models that predict age from molecular data. However, DNA methylation clocks are relatively difficult to interpret and remain distant from gene regulatory networks, a gap that can be complemented by clocks built from another epigenetic layer: chromatin accessibility profiled by ATAC-seq. Existing chromatin accessibility clocks predict age from bulk ATAC-seq data, thereby averaging over the epigenetic heterogeneity across cells that drives aging. We hypothesized that a chromatin accessibility clock trained at the level of individual cell types, using pseudobulk profiles derived from single-nucleus ATAC-seq (snATAC-seq) data, would be particularly useful for characterizing cell type-specific aging. We focused on the brain, a highly heterogeneous tissue whose diverse cell types age asynchronously, and assessed how well cell type-specific accessibility clocks can predict chronological age, capture rejuvenation from genetic perturbation, and detect age acceleration in age-associated neurodegenerative disease. To this end, we introduce a set of cell type-specific and all-cell aging clocks built from snATAC-seq profiles of the prefrontal cortex (PFC) of 357 human donors (15 to 100 years), which generalize to accurately predict age across brain regions and species. Beyond healthy aging, these PFC clocks captured the rejuvenating effects of SIRT6 overexpression in mouse liver and cell type-specific age acceleration in Alzheimer’s disease (AD) and Parkinson’s disease, with microglial age acceleration correlating most strongly with pathology among major cell types, and with female oligodendrocytes and OPCs showing the largest sex differences in age acceleration. Interpreting the clocks further revealed the regulatory elements, genes, pathways, and motifs underlying these signals across species, disease, and perturbation, including repression of the NF-*κ*B pathway in SIRT6 transgenic mice, upregulation of immune and inflammatory pathways in severe AD, and conserved age-predictive peaks related to histone regulation, metabolism, and neuronal survival across brain regions and species. Together, these results establish PFC snATAC-seq aging clocks as a generalizable tool that accurately predicts age and captures cell type-specific perturbation effects of rejuvenation and disease on the epigenetic landscape, providing both a means to evaluate perturbations and insight into the epigenetic mechanisms of aging and disease.

## 1 Introduction

Aging is a major risk factor for many chronic diseases, including cardiovascular disease, cancer, metabolic disorders, and neurodegeneration.^1,2^ Alzheimer’s disease (AD), the most common form of dementia, exemplifies this dependency: its incidence roughly doubles every five years after age 65,^3,4^ yet only a small fraction of cases can be explained by germline genetics, leaving aging itself as the primary risk factor. Understanding the molecular drivers of aging is therefore essential both for extending healthy lifespan and for identifying the cellular and regulatory programs that, when destabilized, predispose individuals to age-associated disease.

Among the proposed drivers of aging, dysregulation of the epigenome has emerged as an important mechanism.^2^ The information theory of aging hypothesizes that aging is driven by the progressive accumulation of epigenetic noise and the gradual loss of the epigenetic information that maintains cellular identity.^5^ Consistent with this view, the epigenome drifts with age in both post-mitotic and proliferating stem cells: changes in DNA methylation, histone modifications, and chromatin accessibility progressively destabilize gene regulatory programs and erode cell-type identity, an effect further magnified in age-associated disease.^6^ This conceptual framework motivates studying aging through the lens of the epigenome.

Because aging is a gradual and continuous process, it is amenable to regression-based molecular models that predict age from measurable features. Such models, termed “aging clocks,” take a high-dimensional molecular profile as input and return an estimate of chronological age as output. The residual between the clock’s output (“biological age”) and chronological age serves as a measure of accelerated or decelerated aging.^7,8^ Aging clocks have been built from a range of molecular modalities, including bulk transcriptomes,^9,10^ metabolomes,^11^ proteomes,^12,13^ and histone modifications,^14^, but clocks built from DNA methylation have stood out for their accuracy and robustness across tissues. Indeed, DNA methylation clocks demonstrate that age can be predicted with remarkable accuracy from a small set of CpG sites in bulk tissue, and that their residuals associate with mortality, disease risk, and lifestyle exposures.^7,8^ However, DNA methylation clocks are typically defined on a fixed and sparse set of CpG sites, are relatively difficult to map onto specific gene regulatory programs, and have been developed mostly at the bulk-tissue level, limiting their cell-type and regulatory interpretability. Chromatin accessibility, profiled by ATAC-seq, complements DNA methylation by surveying the active regulatory genome more broadly: it captures the cis-regulatory elements that orchestrate cell type-specific gene expression and respond to age-associated regulatory rewiring,^15,16^ offering a more direct readout of the gene regulatory programs that underlie aging.

A further hallmark of aging is its heterogeneity: cells, tissues, and individuals age asynchronously and along distinct trajectories.^17^ The aging brain epitomizes this complexity, comprising diverse neuronal and glial populations whose molecular profiles, aging trajectories, and disease vulnerabilities differ substantially.^6,18,19^ Conventional bulk profiling, however, aggregates this heterogeneity into a single average signal, obscuring cell type-specific vulnerabilities. Single-cell technologies overcome this limitation by resolving individual cell states, and consequently, single-cell transcriptomic aging clocks have begun to emerge across diverse systems, including the lung,^20^ heart,^21^ peripheral blood and immune compartments,^22,23^ neurogenic regions of the mouse brain,^24,25^ the spatial architecture of the aging brain,^26^ the fruit fly head,^27^ and several pan-tissue resources of biological age.^28–30^

Progress on the single-cell epigenetic side has been more limited. Single-cell DNA methylation has been used to profile epigenetic age in individual cells,^31^ and recent work has identified clock-like chromatin accessibility loci as readouts of mitotic age in snATAC-seq data.^32^ One chronological age clock has been built directly from single-cell chromatin accessibility in blood;^33^ however, to our knowledge no such clock exists at the cell-type level for the brain, leaving the asynchronous epigenetic aging of brain cell types, and its disruption in age-associated neurodegeneration, unresolved.

Here we hypothesize that cell type-specific aging clocks can be built from snATAC-seq profiles of the human prefrontal cortex (PFC) and are generalizable across brain regions and species. We show that clocks trained on cell type-resolved accessible regions from the PFC of 357 human donors generalize in predicting chronological age across cohorts, brain regions (hippocampus and hypothalamus), tissues (retina and liver), and species (mouse and zebrafish), as well as to bulk sequencing resolution. We further demonstrate that these PFC clocks recover the rejuvenating effects of SIRT6 overexpression in mice, as well as age acceleration in neurodegenerative disease in a sex-specific manner. Finally, through model interpretation, we reveal and prioritize the cell type-specific, aging-associated regulatory elements, genes, pathways, and motifs that drive these predictions.^15^ Together, the PFC clocks provide a cell type-resolved readout of epigenetic age that can be used to probe the regulatory mechanisms of aging and to evaluate cell type-specific perturbations, including disease and rejuvenation strategies.

## 2 Results

### 2.1 All-cell and cell type-specific PFC aging clocks accurately predict chronological age across cohorts, brain regions, and species

To facilitate investigation of aging signatures in single-cell ATAC-seq data for the aging brain across cell types, we established a set of clocks at two levels of resolution: a single donor-level (“all-cell”) clock and a set of cell type-specific clocks, one for each major cell type (Fig. 1a). The clocks were first trained and validated on the largest healthy human prefrontal cortex (PFC) snATAC-seq dataset to date, profiling 1.5 million cells with over 521,000 peaks from 357 healthy donors aged 15 to 100 years (the healthy PFC cohort).^34^ Data were preprocessed and peaks were pseudobulked to obtain a donor-by-peak matrix (Methods), which was used to train cross-validated ridge regression aging clocks (hereafter, “PFC clocks”) for all cells and for each cell type.

**Figure 1:**
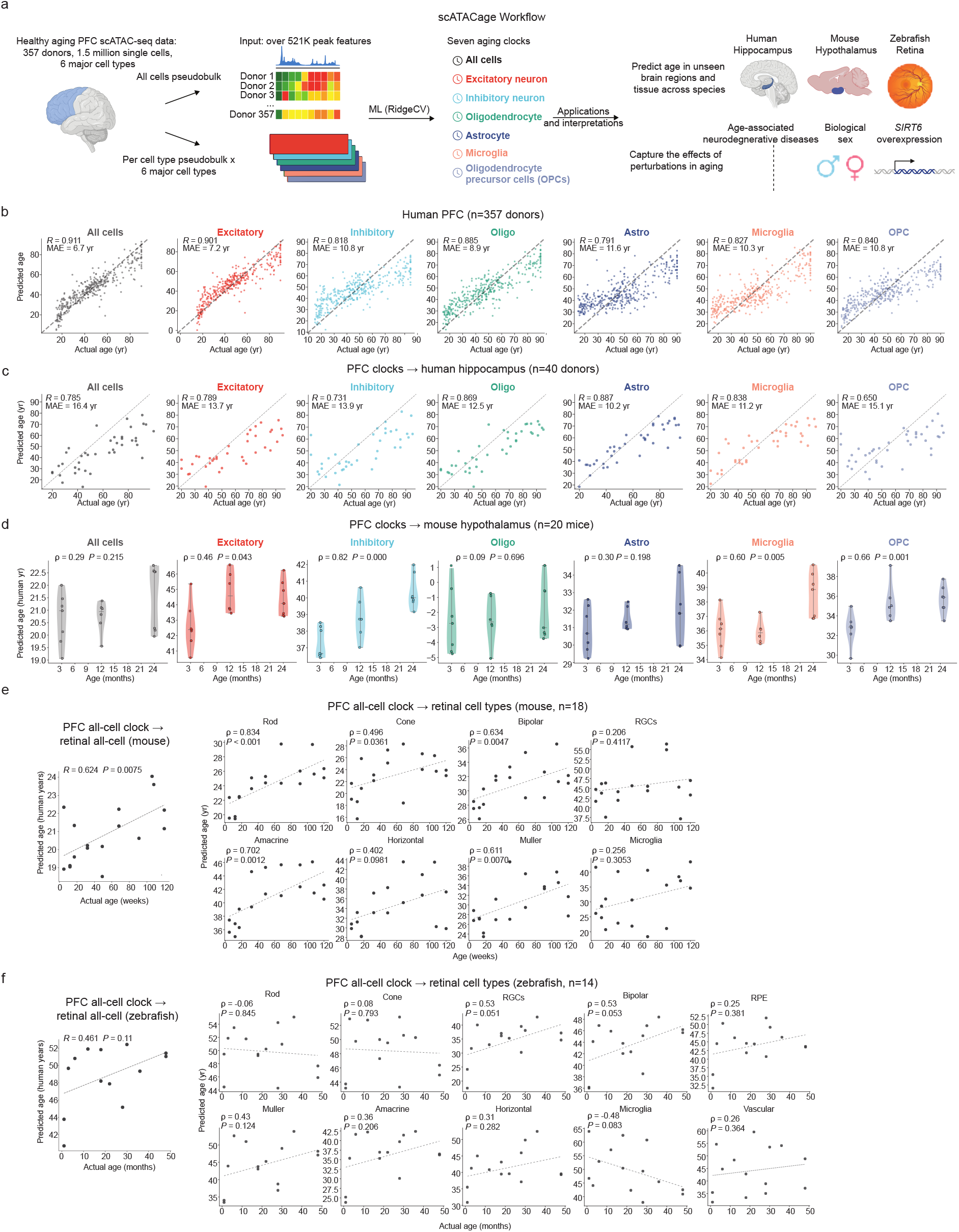
PFC clocks predict chronological age across cohorts, brain regions, and species. **a.** Workflow for training all-cell and cell type-specific pseudobulk models on healthy PFC donors and applying the trained models to cohorts and datasets spanning different brain regions and species. **b-f.** All-cell and cell type-specific clocks accurately predict age, both within the PFC cohort and across brain regions and species.

We assessed the generalizability of the PFC clocks across additional cohorts, brain regions, and species, at both the all-cell and cell type-specific levels (Fig. 1a). First, five-fold cross-validation was performed on the healthy PFC cohort^34^ to assess the clocks’ ability to predict age within a single dataset. To avoid data leakage, the train-test split was performed at the individual level rather than the cell level, with pseudobulks from approximately 71 individuals held out for testing in each fold. After model selection and regularization tuning (Supplementary Figs. S1–S2), the PFC clocks accurately predicted age at both the all-cell and cell type-specific levels (Pearson *r* = 0.791 to 0.911, MAE = 6.7 to 11.6 years; Fig. 1b), despite a relatively moderate sample size (*n* = 357 across the full data) and high feature dimensionality (521,000 features). Performance varied across major cell types: excitatory neurons performed best (*r* = 0.901, MAE = 7.2 years); microglia (*r* = 0.827, MAE = 10.3 years), oligodendrocytes (*r* = 0.885, MAE = 8.9 years), and OPCs (*r* = 0.840, MAE = 10.8 years) came next; and astrocytes performed the worst (*r* = 0.791, MAE = 11.6 years), particularly at younger ages (Fig. 1b). This pattern, with excitatory cells predicting age best and astrocytes worst, is consistent with the differential analysis from the source study of the healthy human PFC cohort, whose authors reported that excitatory neurons had the largest number of age-associated genes across cell types, whereas astrocytes showed no clear age-associated structure in either gene expression or chromatin accessibility.^34^

Next, we examined the PFC clocks’ ability to generalize to an unseen brain region, using a hippocampus cohort of 40 healthy human donors.^35^ We aligned the hippocampus peaks using the PFC coordinates as a reference (Methods), then trained a new set of PFC clocks on the full healthy PFC cohort and applied them to the hippocampus data; for the cell type-specific clocks, cell types in both cohorts were mapped to the six major brain cell types (Methods). The PFC clocks accurately predicted age in the hippocampus (Pearson *r* = 0.650 to 0.887, MAE = 10.2 to 16.4 years; Fig. 1c), suggesting that they capture aging mechanisms shared across brain regions in an independent cohort. Unlike in the healthy PFC cohort, where excitatory neurons performed best, the PFC clocks generalized best to glial cells in the hippocampus, including microglia (Pearson *r* = 0.838, MAE = 11.2 years), oligodendrocytes (Pearson *r* = 0.869, MAE = 12.5 years), and astrocytes (Pearson *r* = 0.887, MAE = 10.2 years), rather than to neurons (Pearson *r* = 0.789, MAE = 13.7 years in excitatory neurons; Pearson *r* = 0.731, MAE = 13.9 years in inhibitory neurons). These results indicate that aging signatures in glial cells may be shared across brain regions, in contrast to neurons, which show pronounced regional specificity and heterogeneity. This result also aligns with the source study for the hippocampus cohort, whose authors noted greater changes in transcriptional state and cell abundance in astrocytes and microglia than in other cell types, such as neurons.^35^

Having validated the generalizability of the PFC clocks across unseen cohorts and brain regions, we next evaluated their performance across both distinct brain regions and species using independent datasets from mouse and zebrafish. First, we analyzed a mouse hypothalamus dataset with 20 mice covering three age groups: young (3 months), middle-aged (12 months), and old (24 months). Unlike the PFC and hippocampus, which support cognition and are relatively less conserved across species, the hypothalamus is an autonomic region that regulates basic physiological processes, including sleep, appetite, and energy homeostasis; consequently, it is more highly conserved across species and shows cell type-specific responses to age, with inflammation in some cell types being amenable to intervention and even lifespan extension.^36–38^ To test whether the PFC clocks can predict age in the mouse hypothalamus, we aligned the PFC and mouse datasets prior to training. Briefly, mouse coordinates (mm10) were lifted over to human coordinates (hg38) and mapped to the PFC peak set, and cell type annotations were harmonized to the six major brain cell types (Methods). Encouragingly, ages predicted by the PFC clocks correlated positively with mouse age across most cell types (Fig. 1d), despite the distinct molecular landscape of the hypothalamus relative to the PFC.^39^ Correlations were strongest in inhibitory neurons (Spearman *ρ* = 0.82) and glia (microglia *ρ* = 0.60, OPCs *ρ* = 0.66), and weakest in oligodendrocytes (*ρ* = 0.09). The strong performance in inhibitory neurons suggests that epigenomic changes in the aging hypothalamus are most pronounced in this population, which may reflect the prominent role of GABAergic dysfunction in hypothalamic aging: the aging hypothalamus develops an excitation–inhibition imbalance driven by GABAergic dysregulation, alongside changes in Sirt1, mTOR, NF-*κ*B, and AMPK signaling,^40^ which may produce a strong and conserved age signal in inhibitory populations. GABAergic hypothalamic neurons, including energy-homeostatic populations of the arcuate and dorsomedial nuclei, have themselves been implicated in systemic aging and lifespan regulation.^37^ By contrast, the weak oligodendrocyte correlation may reflect a limited age signal in hypothalamic oligodendrocytes or poorer cross-species correspondence of oligodendrocyte regulatory elements between the human PFC and mouse hypothalamus, an open question that warrants further investigation.

We next evaluated the PFC clocks on a more challenging dataset: the aging retina from 18 mice and 14 zebrafish. Despite its anatomical distance from the PFC, the retina belongs to the central nervous system (CNS), undergoes age-related degeneration, and represents a system for testing rejuvenation strategies, including epigenetic reprogramming.^41,42^ To align the healthy human PFC and retinal cohorts, we lifted the human PFC (hg38) coordinates to the mouse (mm10) and zebrafish (danRer11) coordinates and mapped the lifted PFC peaks to the mouse and zebrafish peak set.

Retinal cell types differ from those of the brain along several axes. Developmentally, retinal cells arise from the optic cup through a specialized program distinct from most of brain development.^43^ Their neurochemistry is also unusual: most retinal neurons are axon-less interneurons^44^ that can release excitatory neurotransmitters such as glutamate and acetylcholine, as well as the less common inhibitory neurotransmitter glycine, and retinal glial cells form direct electrical couplings with certain neurons.^45^ The eye’s immune environment is also distinct, relying on a specialized lymphatic system for immune privilege.^46^ As a result of these developmental, neurochemical, and immunological differences, retinal cell types are difficult to map precisely onto PFC cell types, whether functionally or molecularly. Given this mismatch, we evaluated the all-cell PFC clock, rather than the cell type-specific clocks, across each retinal cell type; trained on a diverse mixture of cell types, the all-cell clock captures an aging signal that transfers effectively across distinct cellular contexts. Applied to the mouse and zebrafish retinal cohort, the all-cell PFC clock’s predicted ages were positively correlated with mouse retinal age in all cell types (Spearman *ρ* = 0.20 to 0.83; Fig. 1e) and with zebrafish retinal age in seven of nine cell types (Spearman *ρ* = 0.08 to 0.53, excluding rod and microglia; Fig. 1f).

Collectively, these results show that PFC clocks capture conserved aging signatures that generalize across multiple unseen cohorts, distinct brain regions, broader CNS tissues, and species, despite being trained solely on the healthy human PFC without retraining on downstream cohorts. As expected, the performance of the PFC clocks for predicting chronological age on the downstream cohorts decreased as the species and brain region of the downstream cohort diverged further away from the human PFC. Nevertheless, the PFC clocks’ age predictions were positively correlated with the chronological age across all cell types in all cohorts tested, with the exception of the zebrafish retina. This motivates the use of PFC clocks as a quantitative tool to evaluate how therapeutic interventions and age-associated neurodegenerative pathologies modulate biological aging.

### 2.2 PFC clocks predict SIRT6-mediated rejuvenation and identify associated peaks and pathways in bulk ATAC-seq

Motivated by these positive results, we next tested whether the PFC clocks could detect the effects of rejuvenation perturbations, serving as a proof of concept for their utility in intervention screening and drug development. Accordingly, we applied the PFC all-cell clock to a bulk ATAC-seq dataset of 28 male mouse livers, which included 12 mice with an overexpression of SIRT6, a histone deacetylase known to reverse age-associated chromatin changes.^47^ Encouragingly, in both the 14 young (5–7 months) and 14 old (18–21 months) mice, the PFC all-cell clock predicted lower average predicted ages for SIRT6 transgenic mice relative to wild-type controls (Fig. 2a), with the decrease in predicted age significant in old mice (*p = 0.03*, one-sided Mann-Whitney U test). This result demonstrates not only cross-tissue (PFC to liver) and cross-platform (single-cell to bulk) transferability, but crucially, the clock’s capacity to detect biological age reversal.

**Figure 2:**
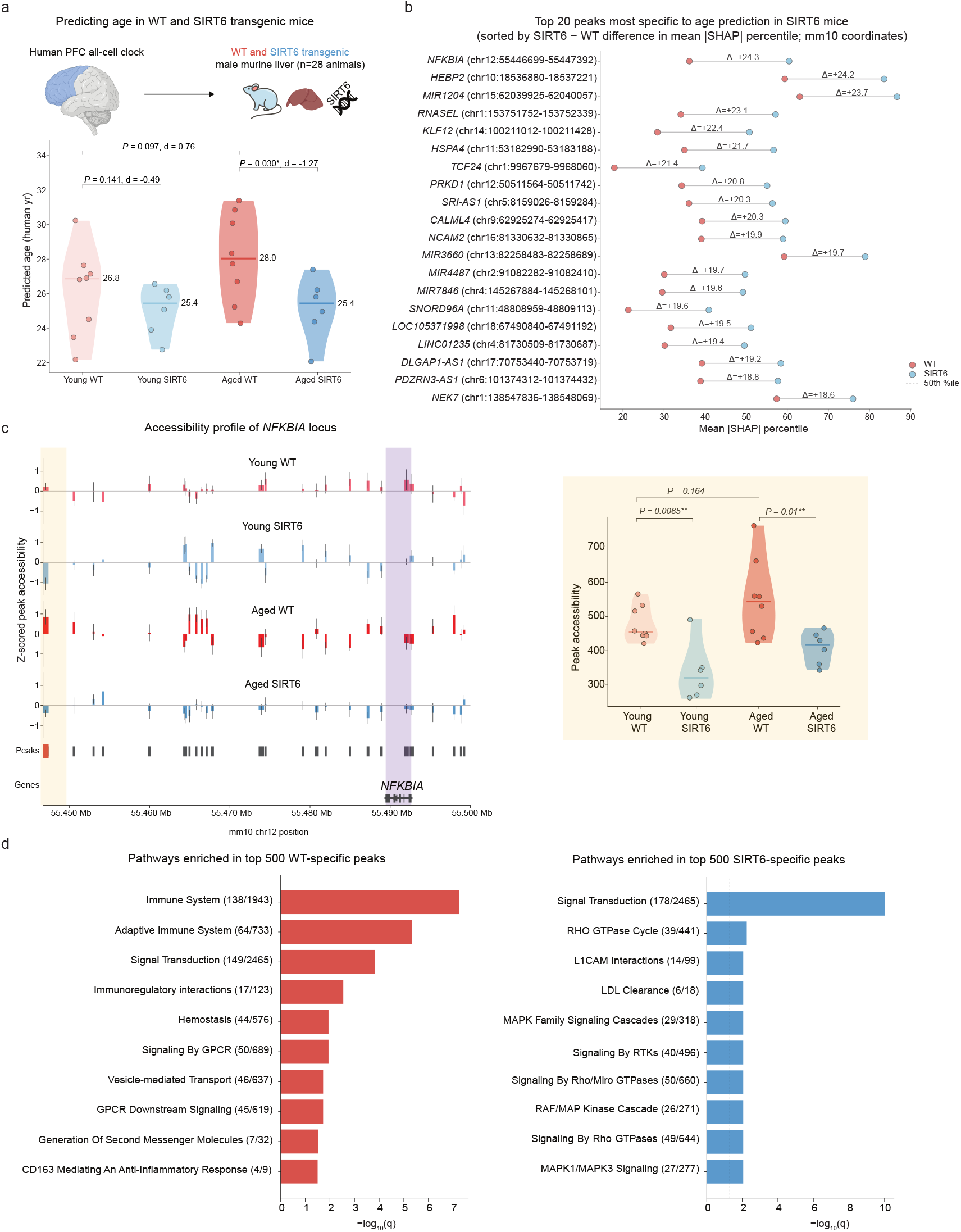
Healthy PFC clocks capture SIRT6-mediated rejuvenation and associated peaks and pathways in male murine liver bulk ATAC-seq. **a.** In both young and old mice, the all-cell healthy human PFC clock predicts lower mean age in SIRT6-overexpressing transgenic mice than wild-type mice, with significance in aged mice (*p = 0.03*; one-sided Mann-Whitney U test). Cohen’s *d* effect sizes and one-sided Mann-Whitney U test *p*-values are displayed for the comparison between young and aged mice, and wild-type and SIRT6 transgenic mice. **b.** Peaks (in mm10 coordinates) most specific to predicting age in SIRT6-overexpressing mice, sorted in descending order by the difference in mean absolute SHAP percentile between SIRT6-overexpressing and wild-type mice. The percentile difference is displayed above each line. c. *Left inset:* ATAC-seq accessibility of the *NFKBIA*-adjacent peak increases with age but decreases with SIRT6 overexpression, consistent with SIRT6’s inhibitory effect on the NF-*κ*B pathway. The decrease in *NFKBIA*-adjacent peak accessibility in SIRT6 relative to WT mice is significant in both young (*p = 0.01*; one-sided Mann-Whitney U test) and aged (*p = 0.01*; one-sided Mann-Whitney U test) mice. *Right panel*: Genomic track plot (in mm10 coordinates) of *NFKBIA* and adjacent peaks, showing differential accessibility in response to both age (young vs. aged) and treatment (WT vs SIRT6). **d.** Top 10 pathways most specific for age prediction in WT mice (left) and SIRT6 mice (right). Peaks were ranked by the difference in mean absolute SHAP between SIRT6 and WT mice (and vice versa), and the the top 500 peaks were tested for enrichment using GREAT with the Reactome database. The top 500 WT-specific aging peaks were enriched for inflammatory-response and immune pathways, whereas SIRT6-specific aging peaks were largely signaling pathways. The vertical dashed line corresponds to a false discovery rate (FDR) threshold of 0.05.

We next asked whether clock interpretability could reveal the regulatory features underlying SIRT6-mediated rejuvenation. For this purpose, we used the SHapley Additive exPlanations (SHAP) framework,^48^ which provides a quantitative measure of how important each input feature is relative to a given clock’s prediction. By computing the difference in mean absolute SHAP percentiles between SIRT6 transgenic and wild-type (WT) mice, we prioritized the top age-predictive peaks driving clock predictions in the transgenic cohort (Methods). The top peaks leveraged by the clock were linked to stress- and immune-response genes, including *HEBP2*, which induces necrotic cell death through mitochondrial membrane permeabilization, *RNASEL*, which is involved in innate immunity, and *KLF12*, a regulator of cell growth (Fig. 2b).

Consistent with age-associated activation of NF-*κ*B, chromatin accessibility at the *NFKBIA*-linked locus increased with age; however, accessibility was significantly lower in SIRT6 transgenic mice than in wild-type controls at both ages (one-tailed Mann-Whitney U test; young *p* = 0.01, aged *p* = 0.01; Fig. 2c). We interpret this reduced *NFKBIA* accessibility in SIRT6 mice as a signature of dampened NF-*κ*B activity, consistent with SIRT6’s established role in suppressing NF-*κ*B-mediated stress and inflammatory responses^49^. Together with the clock’s prioritization of this locus, this suggests that the PFC clock captures SIRT6-mediated suppression of NF-*κ*B signaling as a candidate mechanism of biological rejuvenation.

Notably, the top SIRT6-specific peak was mapped nearest to *NFKBIA*, which encodes I*κ*B*α*, an inhibitor of the NF-*κ*B pathway (Fig. 2b). *NFKBIA* is itself a canonical NF-*κ*B target gene, induced as part of a negative-feedback loop, so its accessibility may track NF-*κ*B pathway activity.^50^ Consistent with age-associated activation of NF-*κ*B, chromatin accessibility at the *NFKBIA*-linked locus increased with age; however, accessibility was significantly lower in SIRT6 transgenic mice than in wild-type controls at both ages (one-tailed Mann-Whitney U test; young *p* = 0.01, aged *p* = 0.01; Fig. 2c). We interpret this reduced *NFKBIA* accessibility in SIRT6 mice as a signature of dampened NF-*κ*B activity, consistent with SIRT6’s established role in suppressing NF-*κ*B-mediated stress and inflammatory responses^49^. Together with the clock’s prioritization of this locus, this suggests that the PFC clock captures SIRT6-mediated modulation of NF-*κ*B signaling as a candidate mechanism of biological rejuvenation.

We next evaluated broader pathway enrichment between the two groups. Whereas the top 500 wild-type-specific peaks were dominated by inflammatory and immune-response pathways, the top 500 SIRT6-specific peaks centered on metabolic and homeostatic signaling, including LDL clearance and MAPK signaling cascades (Fig. 2d). This separation aligns with the original transcriptomic profiling of SIRT6 transgenic mice, which demonstrated a shift away from immune pathways toward metabolic regulation.^47^ Taken together, the PFC clock not only predicts reduced biological age in SIRT6 transgenic mice, but also provides interpretability by demonstrating that SIRT6-mediated regulation of NF-*κ*B and downstream stress and immune-response loci serves as core mechanisms of rejuvenation.

### 2.3 PFC clocks identify age acceleration and implicated pathways in age-associated neurodegenerative disease

Beyond characterizing rejuvenation signatures, we asked whether PFC clocks trained on healthy donors could identify accelerated biological aging in the context of neurodegenerative diseases. To this end, the PFC clocks were applied at both the all-cell and cell type-specific levels to predict chronological age in Alzheimer’s disease (SEA-AD; AD) and Parkinson’s disease (PD) cohorts, respectively consisting of 43 and 101 donors.^18,51^ Corrected age residuals were then obtained by computing the difference between predicted and actual chronological age and correcting for regression-to-the-mean artifacts (Methods). The final age residuals were tested for association with disease condition or severity at both all-cell and cell type-specific levels. Excitingly, we observed a positive association between age residuals and both Alzheimer’s disease neuropathologic change (ADNC) level and the presence of Parkinson’s disease (PD) across all cell types (Fig. 3a-b). Among all major cell types, age residuals from microglia, the resident immune cells of the brain, had the highest correlation with ADNC level and the largest effect size for PD presence (Fig. 3c-d).^52^ Given that AD and PD are driven by distinct primary pathologies, this shared positive correlation between disease severity and microglial age residuals suggests that microglial aging represents a common substrate of neurodegenerative vulnerability rather than a disease-specific consequence, aligning with the view of immune responses to aging as a key process associated with neurodegeneration.

**Figure 3:**
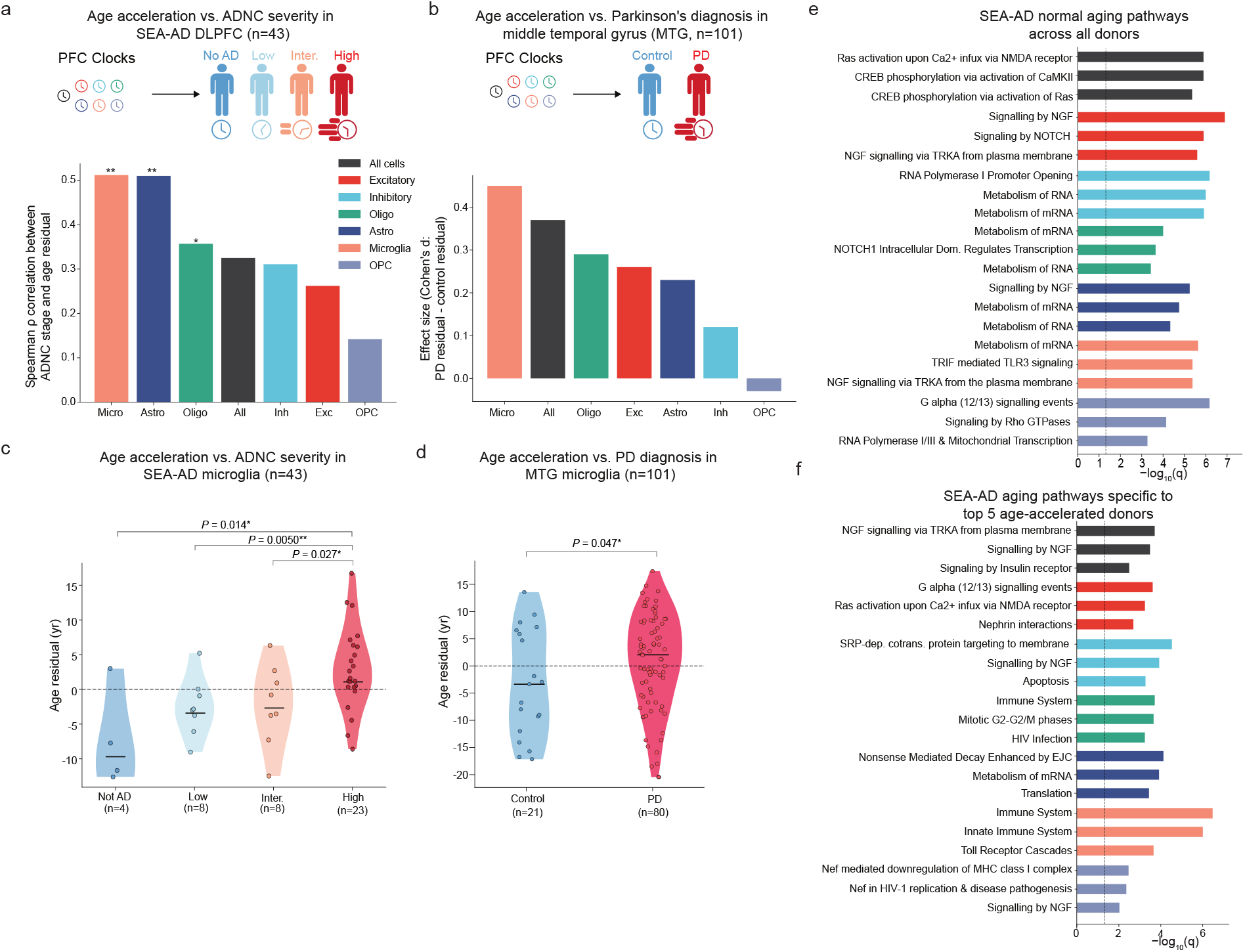
PFC clocks capture age acceleration in neurodegenerative diseases and identify pathways specific to the most age-accelerated donors. **a.** PFC clocks capture age acceleration across major cell types in AD donors (n = 43). In the all-cell clock evaluated on the SEA-AD cohort, ADNC severity is positively correlated with age residual across all cell types, with significance in three cell types by a Spearman’s test, BH-corrected across multiple cell types. (* denotes *q <* 0.05, ** denotes *q <* 0.01, and *** denotes *q <* 0.001). **b.** As in a, for PD donors (n = 101). In six out of seven cell types in PD, the difference in age residual between PD and control donors has a positive effect size. **c.** Age acceleration across ADNC levels in SEA-AD DLPFC microglia. The significance for the increase in age residuals between increasing ADNC severity categories is tested via a one-sided Mann-Whitney U-test. **d.** Age acceleration in PD versus control in Parkinson’s MTG microglia. The significance for the increase in age residuals between the control donors and PD donors is tested via a one-sided Mann-Whitney U-test. **e.** Top SEA-AD aging-associated pathways across all donors. Pathways were obtained using GREAT with the Reactome database, on the top 2000 peaks sorted by mean absolute SHAP across all donors. The vertical dashed line corresponds to an FDR threshold of 0.05. **f.** As in e, for pathways specific to the five most age-accelerated donors. Pathways were obtained using GREAT with the Reactome database, on the top 2000 peaks sorted by the greatest difference in mean absolute SHAP between top five and bottom five donors by corrected age residual.

To pinpoint the molecular processes underlying pathologically accelerated aging rather than normal aging, we hypothesized that pathways specific to highly age-accelerated donors would be more strongly associated with disease pathology than shared aging pathways, exhibiting greater enrichment of immune-response and cell-death cascades. To test this hypothesis, we identified age-predictive peaks across all 43 SEA-AD donors, and separately those most specific to the five most age-accelerated donors (Methods). We then performed genomic region enrichment analysis (GREAT)^53^ on the top 2,000 peaks of each set. The top aging pathways across all SEA-AD donors were dominated by signaling and metabolic programs, such as metabolism of mRNA and RNA, CREB phosphorylation, and signaling by NGF (Fig. 3e). In contrast, the top pathways specific to the most age-accelerated donors were dominated by repair, disease, immune-response, and cell-death programs, such as apoptosis, HIV infection, toll receptor cascades, and nonsense-mediated decay (Fig. 3f). The pathways specific to age-accelerated donors are consistent with AD-induced age acceleration, because deficits in the tau protein in AD are known to be linked to nonsense-mediated decay of mRNA,^54^ toll-like receptors contribute to a microglial neuroinflammatory response,^55^ and apoptosis is a known response to A*β* accumulation and connected to neuronal death.^56^ These results show that the PFC clocks capture disease-associated age acceleration and that the cells driving this acceleration are marked by a shift from metabolic and signaling programs toward immune and cell-death programs, mirroring the inflammatory transition seen in accelerated aging.

### 2.4 PFC clocks identify sex differences in age acceleration across cell types in neurodegenerative disease

We next evaluated whether the PFC clocks are sensitive to documented sex differences in neurodegenerative disease. For example, women with high AD pathology show accelerated markers of brain aging, including greater hippocampal atrophy and steeper cognitive decline, than their male counterparts.^57,58^ We therefore investigated whether the PFC clocks could recover this dimorphism in donors with advanced pathology within the SEA-AD cohort, hypothesizing that female High ADNC donors would display higher age acceleration residuals than male High ADNC donors. We found that female donors exhibited consistently higher age residuals, and thus higher biological age, than their male counterparts at the highest ADNC level across all seven cell type-specific clocks (Fig. 4a). Cohen’s *d* for the female-versus-male difference in age residuals (shown above each split violin) was larger in High ADNC than in Not High ADNC donors in six of seven cell types (Fig. 4a). Interestingly, the greatest sex differences in age residuals among High ADNC donors were in oligodendrocytes and OPCs. These cell types play a sex-specific role in Alzheimer’s disease: female oligodendrocytes and OPCs mount a muted transcriptional response to AD-induced demyelination compared with their male counterparts, leaving them more vulnerable to white matter damage and cognitive decline,^59^ whereas male OPCs mount a robust regenerative response through differentiation and remyelination.^60^ This contrast may explain why the PFC clocks’ median residuals in these cell types were positive (faster aging) in female donors but negative (slower aging) in male donors: the negative residuals in males may reflect active remyelination and differentiation, whereas the positive residuals in females may reflect unmitigated white matter damage.

**Figure 4:**
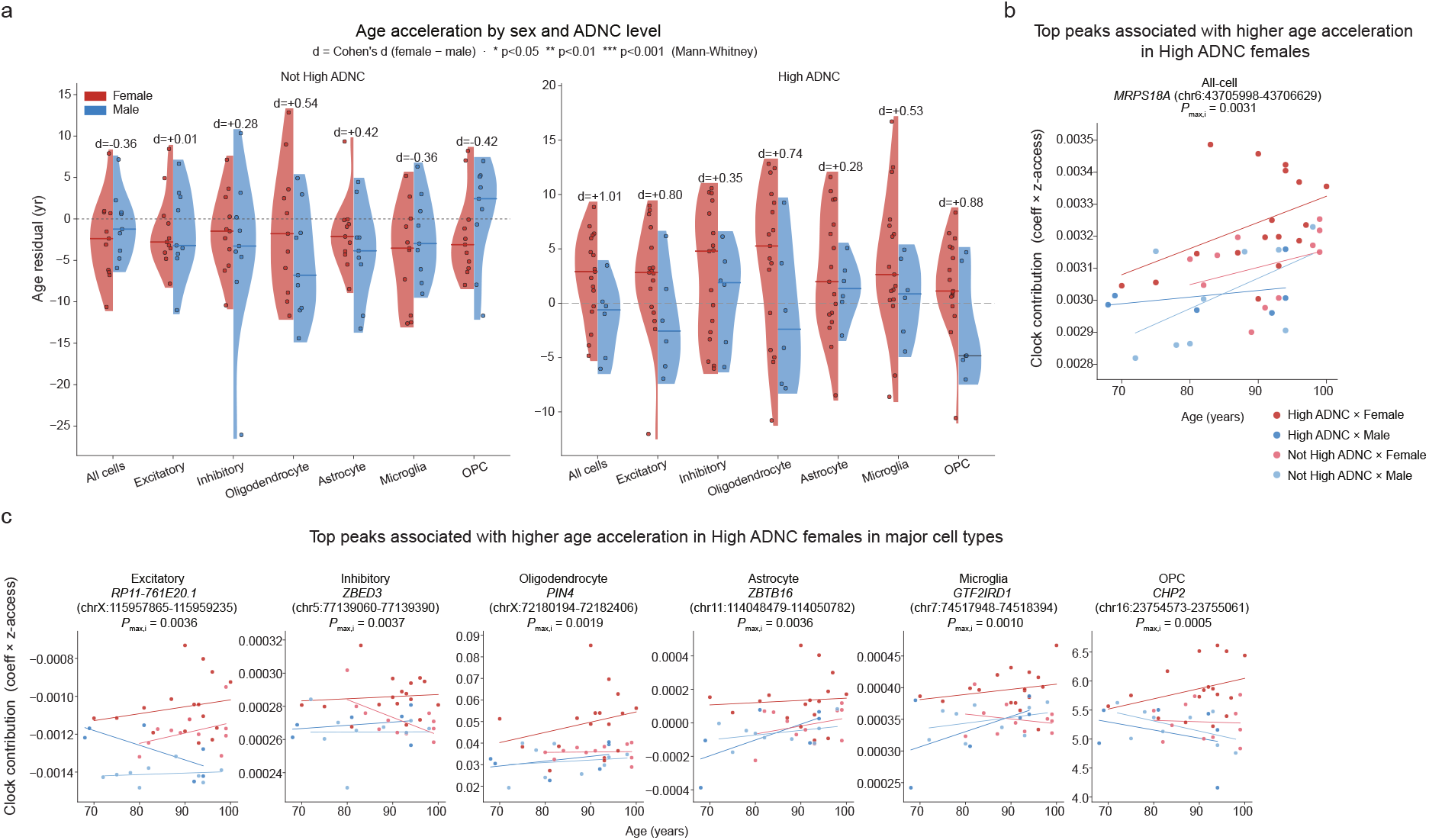
Healthy PFC clocks detect increased age acceleration in females with severe neurodegeneration. **a.** Mean age residuals are greater in female than male donors with high-ADNC AD across all cell types. The effect size (Cohen’s *d*; in bold) of the female-versus-male difference in age residuals is larger for High ADNC than Not High ADNC donors in six out of seven cell types. Due to a limited number of donors in each category, the differences in age residuals in females vs. males, as measured by a one-sided Mann-Whitney U test, was not significant after multiple hypothesis correction across cell types. **b.** Top peaks contributing most to increased age acceleration in High ADNC females. A peak’s contribution to age was defined as its accessibility times its linear model coefficient. For each peak, three one-way ANCOVA tests were conducted, testing for larger peak contribution (adjusted for chronological age) in High ADNC/Female donors versus each of Not High ADNC/Male, Not High ADNC/Female, and High ADNC/Male. Peaks were sorted by the least significant of the three test results, *pmax,i*, as shown in the plots (Methods). Due to the large number of peaks and limited number of donors per category, the top peaks did not achieve significance after correcting for multiple hypothesis testing across all peaks and cell types. Top peaks such as *MRPS18A, PIN4, ZBTB16* belong to gene families with known female-specific responses to severe AD, such as inhibition of autophagy and mitochondrial dysfunction.

To uncover the molecular drivers of this sex-specific vulnerability in severe AD, we analyzed High ADNC donors from the SEA-AD cohort to prioritize peaks driving accelerated aging in females with advanced pathology. Specifically, for each donor and cell type, we computed each peak’s contribution to age as the product of the peak’s clock coefficient and its pseudobulked accessibility. For each peak across cell types, we split donors into four groups (17 High ADNC females, 11 Not High ADNC females, 6 High ADNC males, and 9 Not High ADNC males) and used a set of ANCOVA tests to assess whether a peak’s contribution to age was greater in High ADNC females than in the other three groups, ranking peaks by the significance of this contrast (Methods).

The top all-cell peak was associated with *MRPS18A*, a mitochondrial ribosomal gene linked to cognitive aging in females (Fig. 4b).^61^ Mitochondrial ribosomal genes are strongly tied to aging, with expression-reducing mutations extending lifespan in both *C. elegans* and mice,^62^ and related genes such as *MRPS33* show sex-differential associations with AD cognitive decline.^63^

Interestingly, in oligodendrocytes, an X chromosome-linked peak associated with *PIN4* was most specific to female High ADNC age acceleration (Fig. 4c). Its co-regulated paralog, *PIN1*, regulates tau phosphorylation in AD and mediates apoptosis in adult male oligodendrocytes.^64^ While *PIN1* expression decreases significantly with age across brain regions in both sexes, its correlation with clincal and neuropathological indicators of AD, such as global cognitive function and neurofibrillary tangle density, is significant only in females.^65^ Notably, *PIN1* was also more strongly reduced in severe female AD donors than in female donors with mild cognitive impairment, relative to healthy controls (*p* = 3 × 10*^−^*^4^ versus *p* = 0.027).^65^ Together, these findings suggest that PIN family members, including *PIN1* and *PIN4*, exhibit distinct, sex-dependent responses to severe neurodegenerative pathology.

In astrocytes, the top peak for female High ADNC age acceleration was associated with *ZBTB16*, which is linked to A*β*-induced neurotoxicity and inhibition of autophagy (Fig. 4c). Interestingly, an mGluR5 agonist targeting the ZBTB16-mediated autophagy response mitigated A*β* pathology and reversed cognitive decline in male but not female mice,^66^ suggesting that *ZBTB16* may contribute to female-specific vulnerability to AD-driven age acceleration. Taken together, these findings demonstrate that the PFC clocks capture female-specific age acceleration in severe AD, highlighting candidate regulatory loci—including *PIN4* in oligodendrocytes, *ZBTB16* in astrocytes, and *MRPS18A* at the all-cell level—that converge on tau regulation, autophagic responses, and mitochondrial control.

### 2.5 Interpretation of PFC clocks reveals cell type-specific age-associated regulatory elements and pathways

Having evaluated and interpreted the PFC clocks in the contexts of rejuvenation and pathology, we next sought to uncover the molecular mechanisms underlying clock predictions during healthy aging. To this end, we analyzed age-associated regulatory elements, transcription factor motifs, and pathways using the healthy PFC cohort. For each of the all-cell and cell type-specific PFC clocks, we ranked peaks and their linked genes by their global SHAP value across all PFC donors (Methods).

The top excitatory peak was linked to *POLR2J2*, which encodes a subunit of the RNA polymerase II complex (Fig. 5a) and has been implicated in the aging process across species as well as a potential target for rejuvenation.^68^ The top inhibitory peak was linked to *CD300LB*, a member of the CD300 glycoprotein family that regulates immune processes and is a potential marker of inflammaging (Fig. 5a).^69^ The top oligodendrocyte and astrocyte peak was linked to *GSTM1*, which regulates TNF-*α*-dependent astrocyte responses to inflammation and shows increased expression in astrocytes of the aging mouse frontal cortex (Fig. 5a).^70^ These results suggest that the PFC clocks draw most strongly on peaks related to inflammaging and transcriptional processes when predicting age, which was the feature prioritized in the SIRT6 rejuvenation.

**Figure 5:**
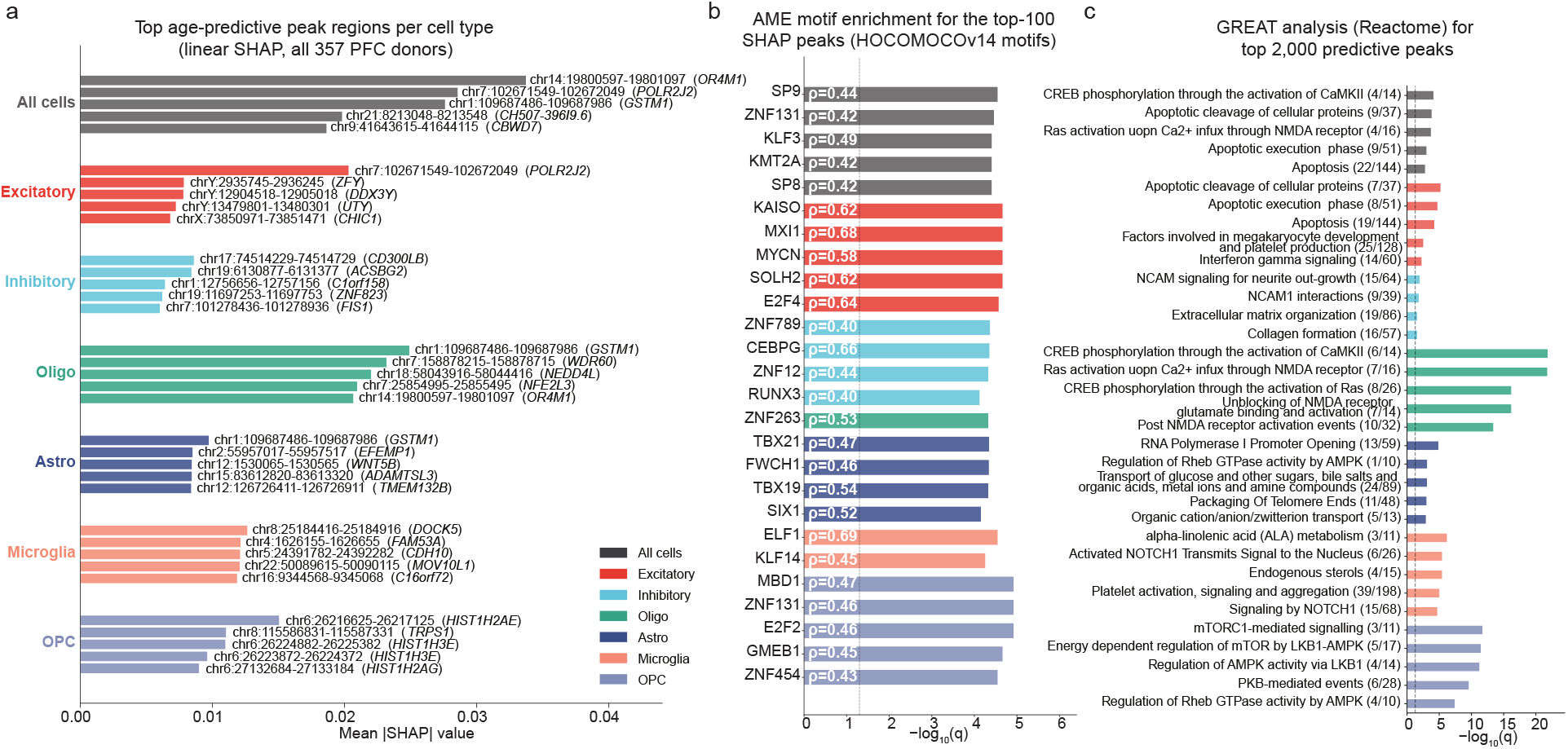
Healthy human PFC clocks identify cell type-specific peaks, motifs, and pathways associated with aging. **a.** Mean absolute SHAP bar plot of all-cell and cell type-specific peaks that contribute the most to age prediction, from PFC samples of 357 human donors. For each cell type, the top three peaks are listed, each with its nearest gene. **b.** Top motifs enriched in the top 100 age-predictive peaks for all cells and per cell type. The three most significant motifs per cell type are listed. Motif significance was obtained using AME^67^ with a Spearman test between a peak’s motif PWM score and the peak’s SHAP importance. The vertical dashed line corresponds to an FDR threshold of 0.05, and the Spearman coefficient *ρ* is shown in the bar to the right of the motif name. **c.** GREAT Reactome pathways enriched in the top 2,000 age-predictive peaks for all-cell and cell type-specific PFC clocks. The three most significant terms per cell type are listed, along with the ratio of the overlap size to the pathway’s gene-set size.

We next examined the transcription factor motifs enriched within the top age-predictive peaks. In inhibitory neurons, the PFC inhibitory clock identified RUNX3 as an important motif for aging (Fig. 5b). The RUNX family (RUNX1, RUNX2, RUNX3) has been extensively studied in aging: RUNX1 in T-cell senescence and inflammaging,^71^ RUNX2 in skeletal aging,^72^ and RUNX3 in the shift toward enhanced granulopoiesis and diminished erythropoiesis with age.^73^ RUNX1 was recently identified as a core master regulator and candidate rejuvenation target in aged human T cells.^71^

In microglia, the PFC clock identified ELF1, a transcription factor that promotes the formation of NLRP3-positive microglia (Fig. 5b). When activated, the NLRP3 inflammasome triggers severe brain inflammation and is linked to accelerated aging and neurodegeneration.^74^

In astrocytes, TBX21 was the top-ranked transcription factor (Fig. 5b). TBX21 is a master regulator of the type I immune response in late-onset AD, is elevated with both age and late-onset AD incidence,^75^ and mediates astrocyte-T cell interactions during neuroinflammation.^76^ Paralleling the top age-predictive peaks enriched for inflammatory pathways, key transcription factors in human PFC aging are strongly linked to inflammaging and neuroinflammation.

Next, for all-cell and per cell type models, we performed a GREAT analysis using the Reactome database^77^ on the top 2,000 age-predictive peaks, correcting for multiple testing across cell types. In OPCs, this analysis identified the mTOR (mechanistic target of rapamycin) and AMPK (AMP-activated protein kinase) signaling pathways, both master regulators of energy and metabolism (Fig. 5c). The mTOR pathway is extensively studied as a central regulator of aging through its roles in metabolism, autophagy, catabolism, and cell proliferation and migration;^78^ its dysregulation is linked to accelerated aging, and mTOR inhibitors such as rapamycin are among the most promising pro-longevity interventions.^79^ Pathways related to the CREB (cAMP response element-binding protein) transcription factor were identified as important for age prediction in both all cells and oligodendrocytes. CREB modulates cognition and cellular function, and reduced CREB activation is associated with aging.^80,81^ In mice, CREB activation in OPCs and oligodendrocytes via cilostazol, a PDE III inhibitor, protects OPCs by promoting white matter repair.^82^ Relatedly, the NMDA (N-methyl-D-aspartate) receptor, implicated in the top pathways for all-cell and oligodendrocytes (Fig. 5c), regulates oligodendrocyte plasticity and myelin maintenance, and NMDA receptor blockade accelerates the deterioration of white matter function with age.^83^ In inhibitory neurons, NCAM (neural cell adhesion molecule) neural signaling pathways were identified as important for predicting age (Fig. 5c). NCAM is depleted in aged mice relative to young mice (roughly 50% lower at 30 months versus 3 months), and its loss is linked to altered synaptic function and architecture.^84^ Together, these findings show that the PFC clocks rely on core signaling, metabolic, and repair pathways, including mTOR, CREB, and NCAM signaling, many of which represent active targets for anti-aging and rejuvenation therapies.

### 2.6 Interpretation of PFC clocks reveals chromatin aging signatures conserved across brain regions and species

Motivated by the PFC clocks’ ability to generalize across cohorts, brain regions, and species, we asked whether the peaks most predictive of healthy PFC age vary with age in similar ways in other brain regions and organisms. To this end, we selected the top 15 PFC-predictive peaks and computed the Spearman correlation between pseudobulked peak accessibility and donor age (the peak-to-age correlation) in the healthy PFC, human hippocampus, mouse hypothalamus, and zebrafish retina datasets. To quantify cross-cohort agreement in how each peak tracked age, we treated the 15 peak-to-age correlations from each cohort as a vector and computed the Spearman correlation between the PFC vector and the vector from each other cohort. We also counted the peaks whose peak-to-age correlation had the same sign (both positive or both negative) in the PFC and each other cohort. The top peak-to-age correlations in the healthy human PFC cohort were positively correlated with those in the human hippocampus, mouse hypothalamus, and mouse retinal datasets (Fig. 6a), and the majority of the top peaks varied with age in the same direction between the PFC and the other datasets (Fig. 6a). Notably, both the cross-cohort correlation and the fraction of direction-concordant peaks decreased monotonically as the target cohort diverged from the human PFC in species and brain region (Fig. 6a). These observations suggest that the peaks most predictive of human PFC age are broadly conserved across species and brain regions, with conservation weakening as evolutionary and anatomical distance increase. This decay of conservation with evolutionary and anatomical distance indicates that a subset of chromatin aging signals are conserved features of the vertebrate regulatory genome, while others are lineage- or region-specific, offering a way to separate universal from context-dependent mechanisms of aging.

**Figure 6:**
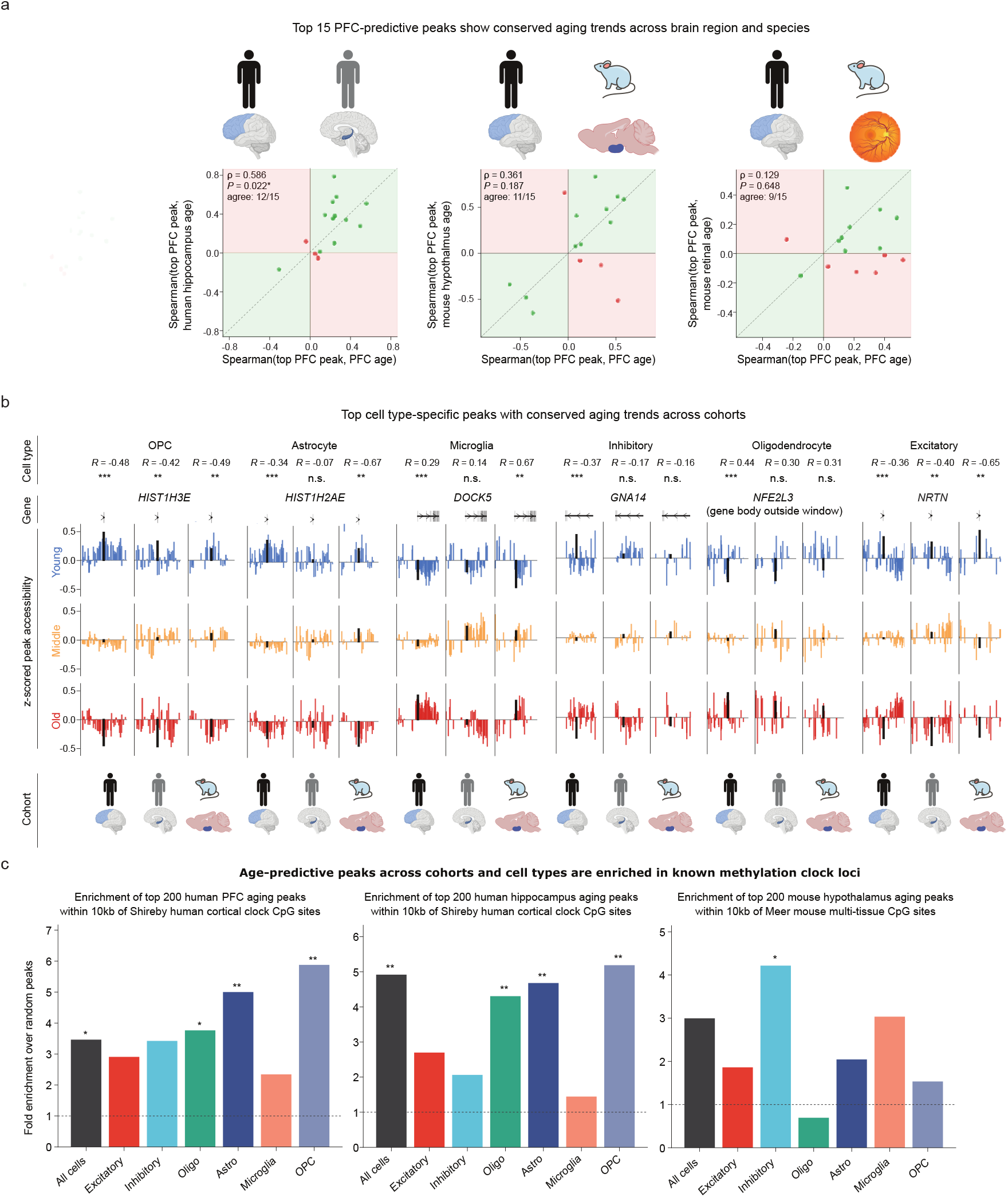
Human PFC clocks identify conserved aging patterns across brain region, species, and modalities. **a.** Relationship (Spearman correlation and sign concordance) between the peak-to-age correlation in healthy PFC and the peak-to-age correlation in each other cohort, across the top 15 PFC-predictive peaks also present in that cohort. **b.** Top PFC-predictive peaks with conserved aging patterns across cohorts (healthy human PFC, human hippocampus, mouse hypothalamus). For each cell type, the top 10 PFC-predictive peaks with the largest absolute correlation summed across the three cohorts were selected. The accessibility profile (log CPM, z-scored across donor pseudobulks) of these selected peaks *±* a 150kb window was plotted across young (youngest third in cohort), middle, and old (oldest third) donors. The black bar in each subplot denotes the selected peak. Asterisks denote significance by a Pearson correlation between peak accessibility and chronological age (* denotes *p <* 0.05, ** denotes *p <* 0.01, and *** denotes *p <* 0.001). **c.** In all three cohorts, the top age-predictive peaks are enriched in known methylation clock loci relative to randomly sampled peaks. A PFC clock peak is considered to overlap a CpG methylation site if their midpoints are within 10 kb of each other. Asterisks denote significant enrichment by a hypergeometric test (* denotes *q <* 0.05, and ** denotes *q <* 0.01).

Given that the top PFC peaks from the all-cell model displayed conserved aging patterns across cohorts, we next sought to identify peaks in the cell type–specific clocks that showed consistent age responses. We used the three datasets in which all six major brain cell types were present (healthy human PFC, human hippocampus, and mouse hypothalamus) and aligned the peaks across all three using PFC peak coordinates as a reference. Among the top ten PFC-predictive peaks for each cell type, we identified the peak with the largest summed absolute peak-to-age correlation across the three cohorts. In excitatory neurons, we found a peak linked to *NRTN* that was consistently downregulated across all three cohorts (Fig. 6b). *NRTN* encodes neurturin, which regulates neuronal survival; AAV-*NRTN* gene therapy has shown neuroprotective and neurotrophic effects in rat and primate models of Parkinson’s disease, and AAV2-*NRTN* is in clinical trials for Parkinson’s.^85,86^ In astrocytes and microglia, histone-related peaks linked to *HIST1H2AE* and *HIST1H3E* showed the most conserved aging patterns (Fig. 6b), both decreasing in accessibility with age (Fig. 6b). *HIST1H2AE* and *HIST1H3E* encode members of the H2A and H3 histone families and are linked to aging-associated histone loss in both human and yeast cells;^87^ because this pattern is conserved across species, such histones have been used to build cross-species histone-modification aging clocks.^88^ Among the top ten microglia peaks, a *DOCK5*-linked peak showed the most conserved aging pattern, consistently upregulated across the three cohorts (Fig. 6b). *DOCK5* is associated with energy balance, and its deletion is connected to dysregulated glucose metabolism and obesity.^89^ *DOCK5* was also the peak with the most significant age-associated accessibility change in microglia in the original PFC study;^34^ our work extends this result by establishing its conserved role across brain regions and species. Taken together, these results show that the PFC clocks recover cell type-specific peaks with conserved aging patterns across brain regions and species, converging on shared aging processes related to metabolism, histone regulation, and neuronal survival.

### 2.7 Age-predictive peaks overlap known methylation clock loci

Having identified peaks with conserved aging patterns across cohorts, we next asked whether these peaks connect to established methylation clocks. Because DNA methylation and chromatin accessibility are closely intertwined, co-regulated processes,^90^ we reasoned that the top peaks for predicting age in the PFC, hippocampus, and mouse hypothalamus cohorts would lie near age-associated CpG loci used by methylation clocks trained on similar brain regions and species.

To this end, we took the top 200 peaks per cell type for predicting age in each cohort. We assessed whether the midpoints of these peaks fell within 10 kb of 347 age-predictive CpG loci from a human cortex clock,^91^ for the clocks evaluated on healthy human PFC and human hippocampus, and of 435 age-predictive loci from a multi-tissue mouse clock,^92^ for the clock evaluated on mouse hypothalamus. In all three cohorts, the top age-predictive peaks showed significant overlap with CpG methylation loci in at least one cell type (In human PFC: *q = 0.002* OPC, In human hippocampus, *q = 0.003* OPC, In mouse hypothalamus, *q = 0.04* Excitatory; BH-corrected one-sided hypergeometric test; Fig. 6c). For the clocks evaluated on healthy human PFC and human hippocampus, the top 200 peaks were significantly enriched in Shireby CpG loci in four of seven cell types (In human PFC: *q = 0.002* OPC, *q = 0.003* Astrocytes, *q = 0.03* Oligo, *q = 0.05* All cells; In human hippocampus: *q = 0.003* OPC, *q = 0.003* Astrocytes, *q = 0.005* Oligo, *q = 0.003* All cells; BH-corrected one-sided hypergeometric test), with positive fold enrichment over randomly sampled peaks in all seven. For the clock evaluated on mouse hypothalamus, the top 200 peaks were significantly enriched in Meer CpG loci in excitatory neurons (*q = 0.04*; BH-corrected one-sided hypergeometric test), with positive fold enrichment over random peaks in six of seven cell types (all but oligodendrocytes).

Next, for each cohort, we sorted peaks by the distance of their midpoints to the nearest CpG locus. For both the healthy human PFC and human hippocampus clocks, a *HIST1H3E*-linked peak was most proximal to Shireby loci in astrocytes, microglia, oligodendrocytes, and all-cells with a distance of 102 bp, and a *HIST1H2AK*-linked peak was most proximal in OPCs with a distance of 23 bp (Table S7). As noted above, *HIST1H3E* and *HIST1H2AK* encode histone family members (H3.1 and H2A) associated with aging-related histone loss.^87^ The independent convergence of our snATAC-seq clocks and Shireby’s methylation clock on histone-gene loci suggests that both modalities detect aging-associated changes in histone regulation, pointing to it as a shared, cross-modality signature of brain aging. In both the healthy human PFC and hippocampus clocks, a *VGF*-linked peak was closest to Shireby loci in OPCs, with a distance of 62 bp. *VGF* generates peptides involved in memory formation and neuroprotection and has been studied as a therapeutic target in Alzheimer’s disease; administration of the *VGF*-derived peptide TLQP-21 reduced amyloid plaque, microgliosis, and astrogliosis in a mouse model of AD.^93^

In the mouse hypothalamus clock, an *ESR1*-linked peak was most proximal to Meer loci in all cells and in excitatory and inhibitory neurons, with a distance of 356 bp. *ESR1* dysfunction is linked to neuroinflammation and pyroptosis and drives severe AD neurodegeneration.^94^ In astrocytes, a peak linked to *C8B*, a component of the membrane attack complex, was closest to Meer loci with a distance of 180 bp; *C8B* is downregulated across multiple tissues in aged mice^95^ and is associated with age-related macular degeneration.^96^

While many of the top ATAC-seq peaks for predicting age in each cohort were proximal to known methylation clock loci, our clocks also identified important, known age-predictive peaks/loci that the methylation clocks did not capture. Among the top 10 PFC-predictive peaks that were also age-predictive in the human hippocampus and mouse hypothalamus, we identified *DOCK5* as an important regulator of energy balance and metabolism, and *NRTN* as a key factor for neuronal survival with neuroprotective effects against diseases such as Parkinson’s (Fig. 6b). However, the *DOCK5*-linked peak identified in the human PFC, human hippocampus, and mouse hypothalamus was not proximal to any methylation clock loci, with the closest CpG loci in the Shireby cortical clock and the Meer multi-tissue mouse clock being more than 2 Mb away. Furthermore, neither was *NRTN*-linked peak proximal to any methylation clock loci: the closest CpG loci in the Shireby cortical clock and the Meer multi-tissue mouse clock were both more than 1 Mb away.

Taken together, these results suggest that our clocks capture both shared and specific age-related mechanisms with established methylation clocks. Notably, our model captures important loci related to energy balance and neuronal survival that were not captured by the methylation clocks. Yet, as a whole, the top age-predictive peaks identified by our clocks and the CpG loci used by established methylation clocks converge on shared age-related mechanisms, including histone loss and neuroinflammation, indicating that accessibility- and methylation-based clocks read overlapping regulatory programs of brain aging.

## 3 Discussion

Here we demonstrate that snATAC-seq clocks trained on healthy human PFC cells can accurately infer chronological age across multiple brain regions and species, without batch correction; capture biological aging by predicting decelerated age under SIRT6-mediated rejuvenation and accelerated age in AD and PD neurodegeneration; and identify the cell type-specific peaks, pathways, and motifs that underlie both chronological and biological aging. To our knowledge, these are the first snATAC-seq clocks to generalize and identify age-associated changes across multiple brain regions (prefrontal cortex, hippocampus, and hypothalamus), species (human, mouse, and zebrafish), tissues (brain, retina, and liver), diseases (AD and PD), and interventions (SIRT6 overexpression). Beyond this generalizability, our analysis suggests that the top age-associated regulatory elements, including peaks, motifs, and pathways, are largely conserved across brain regions and species, pointing to a common aging program shared across evolutionary scales. In particular, our clocks capture conserved peaks related to energy regulation, metabolism, neuronal survival, and histone modification, cellular programs that are consistent with results from multi-species, multi-tissue clocks trained on transcriptomic or methylomic data.^88,97^

A central strength of the snATAC-seq aging clocks is that they read biological, not merely chronological, age. This is illustrated by its sensitivity to rejuvenation effects: our human PFC all-cell clock captured SIRT6-mediated rejuvenation in male murine liver, predicting lower ages for SIRT6 transgenic mice at both young and old ages. Reassuringly, the clock did so through mechanistically interpretable features, prioritizing peaks related to the NF-*κ*B pathway, such as the NF-*κ*B inhibitor *NFKBIA*, and inflammation-related genes such as *HEBP2*, as differentially important for predicting age in SIRT6 mice, consistent with SIRT6’s known role in negatively regulating NF-*κ*B-mediated stress responses.^49^ At the pathway level, the clock likewise associated aging in SIRT6 transgenic mice with signaling and homeostatic programs, in contrast to the inflammation and immune-response programs associated with aging in WT mice, recapitulating the shift away from inflammaging that SIRT6 is thought to drive. The observation that the human PFC-trained clock shows sensitivity to SIRT6-mediated rejuvenation in the murine liver raises the possibility that SIRT6-mediated rejuvenation may be a conserved process observed across tissues and species. As such, we hope our work motivates further investigation of rejuvenation strategies such as SIRT6 modulation, including wet-lab validation of SIRT6 overexpression’s role in dampening NF-*κ*B-mediated inflammation across tissues and species in larger, controlled cohorts.

The same ability to read biological age extended to disease. Our clocks captured increased age acceleration in two distinct neurodegenerative conditions, Alzheimer’s and Parkinson’s, across most major brain cell types, an observation previously reported for transcriptomic clocks^98^ but not, to our knowledge, for chromatin accessibility clocks. Moving from whole-cohort to donor-resolved analysis, we found in the SEA-AD cohort that the peaks most specific to the most age-accelerated donors were enriched for known high-pathology programs, such as apoptosis, nonsense-mediated decay, and toll receptor cascades.^54–56^ Stratifying donors further by sex and ADNC level, we found age acceleration that was enhanced specifically in females with high ADNC, with the largest sex gaps occurring in oligodendrocytes and OPCs. This pattern is corroborated by prior work reporting a muted response of female OPCs and oligodendrocytes to AD-induced demyelination, in contrast to the robust, protective remyelination and differentiation response seen in male OPCs.^59,60^ Consistent with this, several genes linked to the peaks driving female-specific age acceleration, such as *ZBTB16*, are tied to female-specific vulnerability to A*β* pathology in AD, suggesting that our clocks recover not only that females age faster in severe AD but also candidate regulatory elements underlying this difference.^66^

Taken together, these results show that our clocks, though trained only on healthy human PFC cells, capture aging and its associated regulatory elements across multiple tissues, brain regions, species, diseases, and treatment conditions. We attribute this breadth to two features of the training design. First, the clocks were trained on a large, high-quality cohort spanning a wide age range (1.5 million cells from 357 donors aged 15 to 100 years), learning a stable healthy-aging signal that transferred accurately even to much smaller cohorts, such as the mouse hypothalamus (20 donors) and retina (18 donors). Second, because they were trained exclusively on normal-aging donors, the clocks could isolate deviations attributable to disease (AD and PD) or treatment (SIRT6 transgenesis). Beyond these design choices, the clocks demonstrate strong cross-species transfer, which can be attributed to the strong conservation of chromatin accessibility across species, particularly in the brain.^99^

Given these strengths, we envision our clocks being used to study aging in additional brain regions, species, and conditions. Because these clocks already capture SIRT6-mediated rejuvenation, a natural next step is to test other interventions, such as exercise, parabiosis,^24^ or dietary change,^100^ and to prioritize rejuvenation strategies by their age residuals. Conversely, the same framework could rank candidate age accelerators, such as environmental toxins, chronic stress, smoking, and air pollution.^101,102^ A complementary direction is to dissect how intrinsic and extrinsic factors interact: our finding of enhanced age acceleration in females with high ADNC already suggests that sex and disease severity combine non-additively, and future work could extend this to genotype, drug treatment, or environmental exposure, informing precision aging medicine in which interventions are tailored to an individual’s genetics, lifestyle, and exposures.

We also anticipate improvements to the model itself. Expanding the training set with donors from additional brain regions, tissues, and species could further improve generalization, particularly if the model is trained across these settings simultaneously. Here, owing to the modest sample size, we used a ridge clock, as is common for methylation-based clocks;^103,104^ as more scATAC-seq aging data accumulate, nonlinear models such as EpiAgent or PoissonVI could capture nonlinear aging dynamics.^105,106^

Our study also has several limitations. Harmonizing diverse cohorts into six major cell types, while necessary for cross-cohort comparison, collapses finer distinctions, most notably for the retina, whose cell types map only approximately onto brain categories; our cross-species analyses also rely on coordinate lift over, which degrades with evolutionary distance and may underlie the weaker generalization seen for the most distant cohorts and for cell types such as oligodendrocytes. Finally, our mechanistic interpretations are associational, and establishing causal roles will require targeted perturbation. Together with the extensions above, addressing these limitations will further sharpen cell type-resolved chromatin accessibility clocks as a tool for probing the epigenetic mechanisms of aging and disease.

## 4 Methods

### 4.1 Preprocessing of bulk and snATAC-seq cohorts

We used ATAC-seq data from eight previously published studies. For three datasets—Catching 2026, Nagar 2026, and Yu, Osipov, and Hassell (YOH) 2026—full cell-by-peak matrices were provided as preprocessed h5ad or CSV files by the authors. For Zemke 2024, Adams 2024, Lyu 2025, and Menon 2026, the cell-by-peak matrix was obtained by concatenating per-donor processed h5 or rds cell-by-peak matrices. For Gabitto 2024, we combined donor-level fragment files and obtained cell-by-peak matrices by counting the number of fragments that overlapped with peaks in the training cohort (Catching 2026). We normalized peak counts for each dataset using counts per million mapped reads (CPM) normalization followed by log(*x* + 1) scaling. Then, for each dataset, we constructed two levels of pseudobulk data, all-cell and per-cell-type, by computing the mean peak accessibility across all cells per donor and across cells of each cell type per donor, respectively. Cell type annotations were obtained from the authors except Lyu 2025, where cell types were inferred from author-provided cell type marker genes using the paired RNA-sequencing data. A final per-donor pseudobulk scaling was performed by Z-scoring each donor’s pseudobulk across peaks. Due to the relative sparsity of the Zemke 2024 and Lyu 2025 datasets, we additionally performed 5 kb and 50 kb binning of each dataset, in which we constructed donor-by-tile matrices by summing across peaks within each 5 kb or 50 kb window.

### 4.2 Mapping cell types across cohorts

To enable consistent per-cell-type clocks across datasets with differing annotation schemes, we first harmonized cell type labels across brain cohorts to six major cell types: astrocytes, microglia, excitatory neurons, inhibitory neurons, oligodendrocytes, and oligodendrocyte progenitor cells (OPCs). For three datasets—Catching 2026, Menon 2026, and Gabitto 2024—we used author-provided metadata, which includes cell annotations for the six major brain cell types. For the YOH dataset, the “major” cell type column included astrocytes, oligodendrocytes, NG/OPC, immune, GLU, and GABA; we mapped immune to microglia, GLU to excitatory, and GABA to inhibitory. For the human hippocampus cohort (Zemke 2024), the “subclass” cell type column was mapped as follows: DG, CA1, CA2-CA3, and SUB cells were mapped to excitatory neurons; SST, VIP, PVALB, LAMP5, NR2F2, and chandelier cells were mapped to inhibitory neurons; and macrophage and microglia cells were mapped to microglia. The astrocyte, oligodendrocyte, and OPC sub-classes were already labeled as desired. For the mouse and zebrafish retinal cohort (Lyu 2025), cell type labels were not provided in the snATAC multiome data. We therefore inferred cell type labels using the marker genes for each of the major retinal cell types described in the study. We assigned each cell to the cell type whose marker genes had the greatest mean gene expression, using the paired RNA-seq data. The resulting retinal cell types were microglia, Müller, rod, cone, bipolar, amacrine, retinal ganglion, and horizontal cells. Because of the considerable difference in structure and function in retinal cell types and cell types in other parts of the brain,^107^ we opted to use the all-cell PFC clock to predict age in each of the retinal cell types. The all-cell clock is trained on a mixture of PFC cell types rather than a single cell type, and therefore learns a more global aging signal. The SIRT6 cohort (Nagar 2026) contains bulk ATAC-seq samples of murine liver, so mapping to major brain cell types was not possible.

### 4.3 Training and validation of human PFC aging clocks and their application to test cohorts

Before training and validation, we aligned the training and testing cohorts by using the training peak coordinates as a reference and summing over all test cohort peaks that intersected each training peak feature. We then trained all-cell and per-cell-type aging clocks using the RidgeCV regression model from scikit-learn (version 1.6.1), with a 30-point log-spaced sweep of the regularization parameter *α* from 10*^−^*^2^ to 10^6^ (Supplementary Fig. S2). The optimal *α* was chosen via leave-one-donor-out cross-validation on the training set, using mean squared error (MSE) as the scoring measure. We used RidgeCV because it offers a favorable tradeoff between runtime efficiency and interpretability, as well as for its consistent performance in predicting age in downstream cohorts (Supplementary Fig. S1). To apply the trained clocks in the cross-species experiments, we used pyliftover^108^ to lift human (hg38) genomic coordinates over to mouse (mm10) and zebrafish (danRer11) coordinates. We then applied the trained clocks to the aligned test cohorts to obtain donor-level age predictions, evaluating performance using mean absolute error (MAE) and Pearson’s correlation coefficient.

### 4.4 Inferring age acceleration in diseased individuals using aging clocks

Human PFC aging clocks were trained on healthy donors and used to obtain age predictions in two neurodegenerative disease cohorts containing both healthy and diseased donors: Alzheimer’s disease (AD) and Parkinson’s disease (PD). Raw residuals were first computed as residual_raw_ = predicted age − actual age. A linear fit was then applied to correct for the bias of the residuals against actual age, as done in similar aging clocks,^109^ yielding the corrected residual residual_corrected_ = residual_raw_ − lm(residual_raw_ ∼ actual age). To test whether the corrected residuals capture disease-associated age acceleration, we computed the Spearman correlation coefficient between age acceleration and disease staging (ADNC for AD), as well as Cohen’s *d* for the difference in age acceleration between PD and healthy donors.

### 4.5 Obtaining age- and disease-associated regulatory elements and pathways through model interpretation

We computed attributions for each peak using SHAP’s LinearExplainer,^48^ which represents the importance of peak *i* for donor *k* as *ϕ^k^_i_ = β_i_(x^k^_i_ − E[x_i_])*, where *x^k^_i_* is the peak accessibility and *β_i_* is the ridge regression coefficient for the corresponding peak. Thus, *ϕ^k^_i_* indicates the importance of peak *i* in predicting age specifically for donor *k*. We then computed global attributions across all donors, *rank_i_ = 1/N Σ^N^_k=1_ |ϕ^k^_i_|*, and used these values to rank peaks from most to least important for predicting age across the cohort.

In addition to global attributions, we used SHAP to quantify a peak’s importance for predicting age within a subset of donors, by summing absolute donor-level attributions across that subset of donors. We used this approach to obtain the importance of peak *i* for predicting age specifically in SIRT6 mice or in the most age-accelerated donors.

### 4.6 Identifying aging-associated pathways using GREAT and Reactome

To obtain pathways enriched in top age-predictive peaks, we performed GREAT^53^ genomic region enrichment analysis on the top peaks using the Reactome pathway database. We called the GREAT web server via the *submitGreatJob* function in rGREAT (v3.0) with default parameters, and called *getEnrichmentTables* on the job results with the “MSigDB Pathway” ontology, which contains Reactome pathway results. For healthy PFC and SEA-AD we used 2000 top peaks for all cells and per cell type, and for SIRT6 mice we used 500 top peaks.

### 4.7 Identifying group-specific aging peaks and pathways via differential SHAP analysis

To identify peaks and pathways whose contribution to age prediction differs between two groups of donors, we applied a differential SHAP procedure across two pairs of groups: age-accelerated versus age-decelerated donors in SEA-AD, and SIRT6-overexpressing versus wild-type mice.

For each pair of groups, we first computed the mean absolute SHAP percentile for each peak separately within the two donor groups, for all cells, and per cell type. We then defined group-specific peaks as the top peaks with the greatest difference in mean absolute SHAP percentile between the two groups. Finally, we performed a GREAT analysis on these group-specific peaks, yielding group-specific aging pathways for all cells and per cell type.

The two pairs of groups were defined as follows. For SEA-AD age acceleration, the two groups were the five donors with the greatest age acceleration and the five donors with the least age acceleration, yielding age acceleration-specific peaks and pathways. For SIRT6-mediated rejuvenation, the two groups were SIRT6-overexpressing transgenic and wild-type male murine liver samples, yielding SIRT6 transgenesis-specific and wild type-specific aging peaks and pathways.

### 4.8 Identifying peaks and pathways with conserved aging patterns across brain regions and species

To test whether the top age-predictive peaks in the healthy PFC show conserved aging patterns elsewhere, we sorted PFC peaks by mean absolute SHAP value across donors and identified the top 15 PFC-predictive peaks that were also present in the other healthy mammalian cohorts (human hippocampus, mouse hypothalamus, and mouse retina). We did this separately for each testing cohort, using the aligned training and testing peaks used in the clocks in Fig. 1. For each of these peaks, we computed the Spearman correlation between pseudobulked peak accessibility and donor age (the peak-to-age correlation) in the four cohorts. To quantify cross-cohort agreement in how each peak tracked age, we treated the 15 peak-to-age correlations from each cohort as a vector and computed the Spearman correlation between the PFC vector and the vector from each of the other three cohorts.

We then extended this analysis to identify, plot, and interpret specific peaks with conserved aging patterns across three brain regions (human PFC, human hippocampus, and mouse hypothalamus) for all six major brain cell types. For all cells and per cell type, we ranked the top 10 PFC-predictive peaks identified in all three cohorts by the absolute sum of their peak-to-age correlations across the three cohorts, defined peaks with conserved aging patterns as those with the highest absolute sum. Peak coordinates were defined using the PFC cohort, and a PFC peak was considered identified in all three cohorts if it intersected one or more peaks in both the hippocampus and mouse hypothalamus cohorts. To align the three datasets to this common PFC-based coordinate set, counts from peaks in the hippocampus and mouse hypothalamus cohorts that overlapped a given PFC peak were summed.

### 4.9 Motif analysis of top PFC-predictive peaks

For each cell type and all cells, we tested the top 100 mean absolute SHAP peaks for motif enrichment in HOCOMOCOv14,^110^, which contains 1,107 human TFs and 809 mouse ortholog transcription factor (TF) motifs. We used AME^67^ from the MEME suite (v5.5.9)^111^ with max scoring and Spearman testing. AME takes in a ranked list of peaks and, for each motif, computes the Spearman correlation and associated significance between 1) each peak’s mean absolute SHAP score and 2) the odds score of that motif’s position weight matrix (PWM) in the peak sequence.

### 4.10 Identification of peaks associated with accelerated aging in females with High ADNC

Using the 43 SEA-AD DLPFC donors, we stratified donors by ADNC level (High ADNC vs. Not High ADNC) and by sex (Male vs. Female) to form four subgroups. For each peak *i* and donor *k*, we computed the contribution of that peak to the predicted age of that donor, *contrib^k^_i_ = β_i_x^k^_i_*, by multiplying the clock coefficient *β_i_*, by the pseudobulked accessibility *x^k^_i_* of peak *i* in donor *k*. For each peak *i*, we then fit an ANCOVA model across donors,

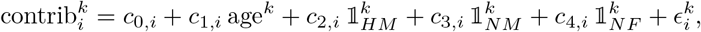

where age*^k^* is the chronological age of donor *k*, the *c_n,i_* are peak-specific ANCOVA coefficients, ε^k^_i_ is the error term for peak *i* in donor *k*, and ‖^k^_HM_, ‖^k^_NF_ are indicator variables denoting whether donor *k* is High ADNC and male, Not High ADNC and male, or Not High ADNC and female, respectively; High ADNC female donors serve as the reference group. We filtered out all peaks with a negative overall contribution to age, that is, peaks with *c*_1*,i*_ *<* 0. We then performed three one-sided tests on the ANCOVA coefficients, for *c*_2*,i*_ *<* 0, *c*_3*,i*_ *<* 0, and *c*_4*,i*_ *<* 0; that is, we tested whether a peak’s contribution to age was reduced in the High ADNC male, Not High ADNC male, and Not High ADNC female groups relative to the High ADNC female group. This yielded three one-sided *p*-values per peak, *p_HM,i_*, *p_NM,i_*, and *p_NF,i_*. Because we required the contribution to age in High ADNC female donors to be significantly higher than in all other subgroups, we ranked peaks by their maximum *p*-value, *p*_max*,i*_ = max(*p_HM,i_, p_NM,i_, p_NF,i_*), with smaller *p*_max*,i*_ ranked higher.

**Table 1:**
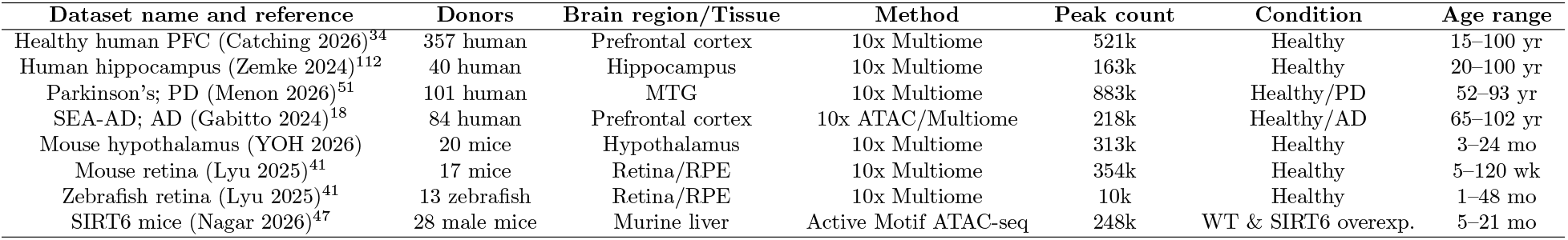
Datasets used to train and evaluate the PFC clocks. We trained and evaluated our PFC clocks on various species, brain regions, and disease conditions. The healthy PFC dataset used to train the model is Catching 2026 (denoted as the first row of the table).

### 4.11 Enrichment of age-predictive peaks in known methylation clock loci

To assess whether age-predictive peaks overlap known methylation clock loci, we took the top 200 peaks per cell type for predicting age in the healthy human PFC, human hippocampus, and mouse hypothalamus cohorts. We assessed their proximity to 347 age-predictive CpG loci from a human cortex clock^91^ for the PFC clocks evaluated on human hippocampus and healthy human PFC, and to 435 age-predictive loci from a multi-tissue mouse clock^92^ for the PFC clocks evaluated on mouse hypothalamus. We considered an snATAC-seq peak proximal to a methylation CpG locus if the midpoint of the peak was within 10kb of the CpG site. Both ATAC-seq peaks and CpG sites tend to be proximal to gene transcription start sites (TSS), and this is especially the case in top age-associated ATAC-seq peaks.^34^ As such, to ensure that the overlap of the ATAC peaks to CpG sites is not confounded by their shared proximity to TSS sites, we generated a set of 200 random peaks where the distance between each peak and its nearest TSS site is the same as the 200 top age-predictive peaks. Specifically, we bucketed the distances between each of the top 200 age-predictive peaks and the nearest TSS into 10 quantiles, and sampled 200 random peaks whose distances with their nearest TSS sites followed the same quantile distribution. We then counted the CpG sites proximal to the top 200 age-predictive peaks, *n*_top_, and those proximal to the 200 TSS distance-matched random peaks, *n*_random_. We then calculated the fold enrichment of the top peaks relative to random peaks, fold enrichment = *n*_top_*/n*_random_. We also conducted a hypergeometric test for the overlap between the top 200 age-predictive peaks and the CpG loci, relative to the 200 random background peaks, with Benjamini-Hochberg (BH) correction across cell types. For each cohort, we plotted the fold enrichment of top versus random peaks as a bar plot across cell types, with the significance of the hypergeometric test indicated by asterisks.

## 5 Data availability

The healthy human PFC data for clock training can be found at https://zenodo.org/records/18394349 (final atac data.h5ad). The human hippocampus multiome h5 files are available at the Gene Expression Omnibus (GEO) under accession ID GSE278576. The mouse retinal multiome h5 files are available under GEO accession ID GSE325478, and the zebrafish retinal multiome h5 files are available under GEO accession ID GSE325620. The male murine liver SIRT6 ATAC-seq data is available under GEO accession ID GSE294103. The Parkinson’s multiome h5 files are available via the Impact of Genomic Variation on Function (IGVF) Data Portal at https://data.igvf.org/. The links for downloading the individual per-donor h5 files is located in Supplementary Table 11 of the accompanying publication, at https://www.biorxiv.org/content/biorxiv/early/2026/03/05/2026.03.05.709922/DC2/embed/media-2.xlsx. The coordinates of the 347 CpG loci of the Shireby human cortical clock can be found under the supplementary data of the accompanying publication, at https://oup.silverchair-cdn.com/oup/backfile/Content_public/Journal/brain/143/12/10.1093_brain_awaa334/5/awaa334_supplementary_data.pdf. The co-ordinates of the 435 CpG loci of the Meer multi-tissue mouse clock can be found under Supplementary File 3 of the accompanying publication, at https://doi.org/10.7554/eLife.40675.022. Peak-to-nearest gene annotations were conducted using pyensembl using ensembl77. The lift over chain files used to lift between human coordinates and mouse/zebrafish coordinates can be obtained at https://hgdownload.soe.ucsc.edu/gbdb/hg38/liftOver/hg38ToMm10.over.chain.gz (for human to mouse), https://hgdownload.soe.ucsc.edu/gbdb/mm10/liftOver/mm10Tohg38.over.chain.gz (for mouse to human), and https://hgdownload.soe.ucsc.edu/gbdb/hg38/liftOver/hg38ToDanRer11.over.chain.gz (for human to zebrafish). The databases used for enrichment analyses are: HOCOMOCOv14 (for motif enrichment), Reactome 2022 (for pathway enrichment). The results published here are in part based on data obtained from the The AD Knowledge Portal, DOI: https://doi.org/10.7303/9618137. The SEA-AD study data were generated from postmortem brain tissue obtained from the University of Washington BioRepository and Integrated Neuropathology (BRaIN) laboratory and Precision Neuropathology Core, which is supported by the NIH grants for the UW Alzheimer’s Disease Research Center (P50AG005136 and P30AG066509) and the Adult Changes in Thought Study (U01AG006781 and U19AG066567). The SEA-AD study is supported by NIA grant U19AG060909.

## 6 Code availability

The code for the training, evaluation, and analysis of PFC clocks can be located under the GitHub repository https://github.com/Noble-Lab/snATAC_aging_clocks.

## Acknowledgements

We thank members of the Noble Lab, as well as Drs. Ran Zhang and Anupama Jha, for their discussions and suggestions for the paper. This work is supported by National Human Genome Research Institute award U01 HG013198 (WSN) and National Institute on Aging F99/K00 award AG083292 (DY).

## 7 Author contributions

PZY and DY contributed equally to this work.

|  | PZY | DY | YX | WSN |
| --- | --- | --- | --- | --- |
| Conceptualization | • | • |  | • |
| Data curation | • | • |  | • |
| Funding acquisition |  | • |  | • |
| Investigation | • | • |  | • |
| Methodology | • | • | • | • |
| Project administration | • | • |  | • |
| Software | • |  |  |  |
| Resources |  |  |  | • |
| Supervision |  |  |  | • |
| Visualization | • | • |  | • |
| Writing—original draft | • | • |  | • |
| Writing—review & editing | • | • | • | • |

## Supplementary information

**Supplementary Fig. S1:**
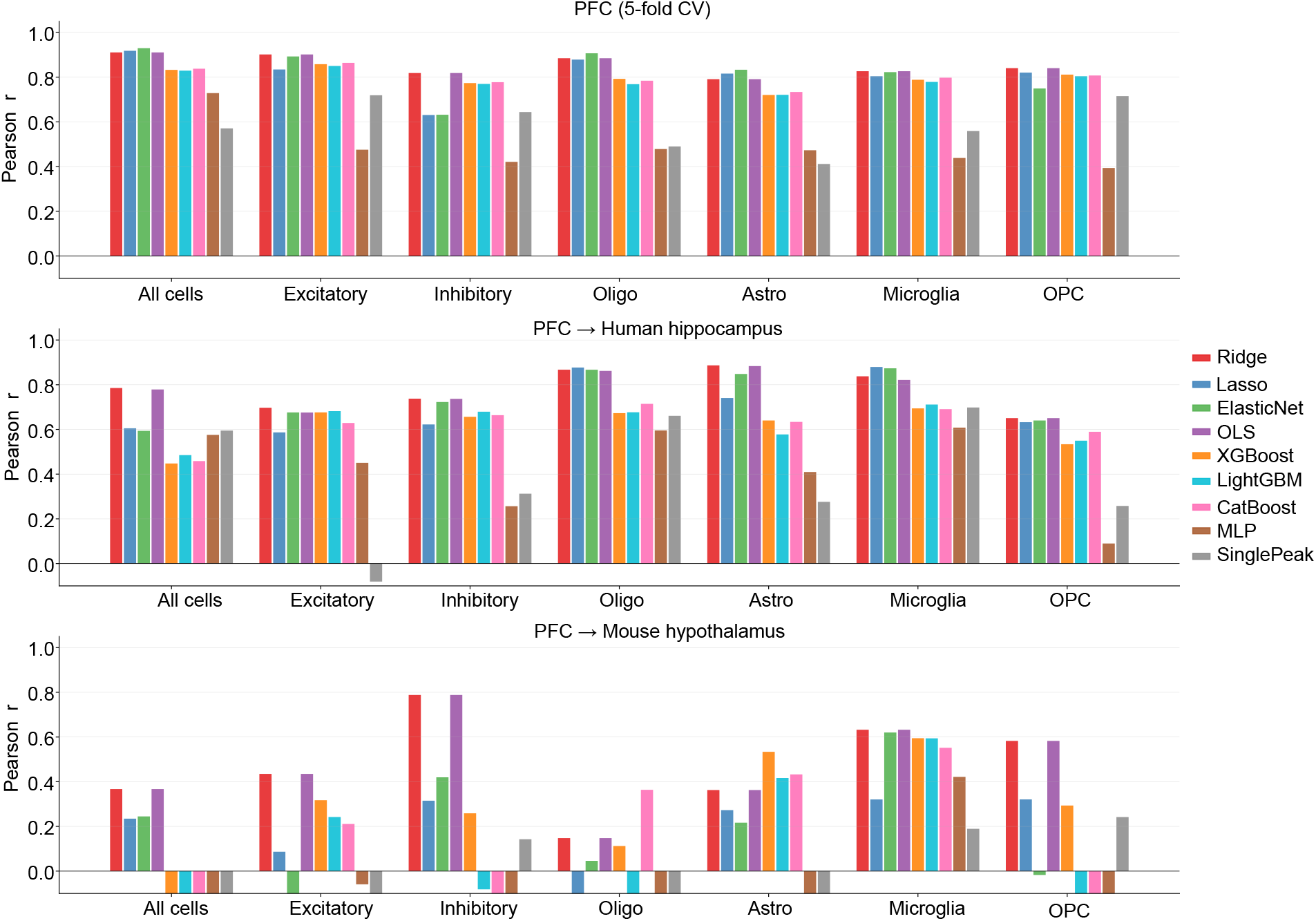
Ridge regression outperforms other modeling formulations in age prediction, including those tested in sc-ChromAging (XGBoost, three-layer MLP, ElasticNet, Lasso, CatBoost, LightGBM, and SinglePeak, ordinary least squares), across most cohort and cell type combinations. The SinglePeak baseline uses the peak most strongly correlated with age in the training set to linearly predict age in the test set.

**Supplementary Fig. S2:**
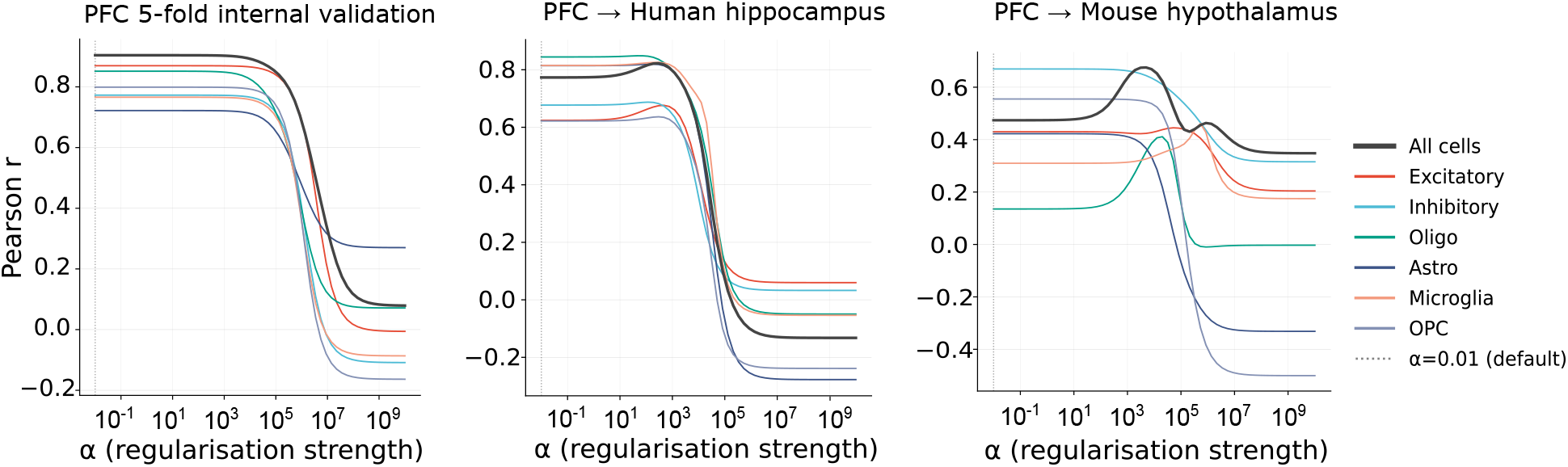
Test performance versus hyperparameter sweep for ridge regression across the three datasets. The optimal regularization strength increases with greater domain shift (from left to right: same-cohort internal validation; different cohort and brain region; different cohort, brain region, and species). However, because all the clocks were trained and validated on healthy PFC, the *α* selected for all downstream tasks is *α* = 0.01.

**Supplementary Fig. S3:**
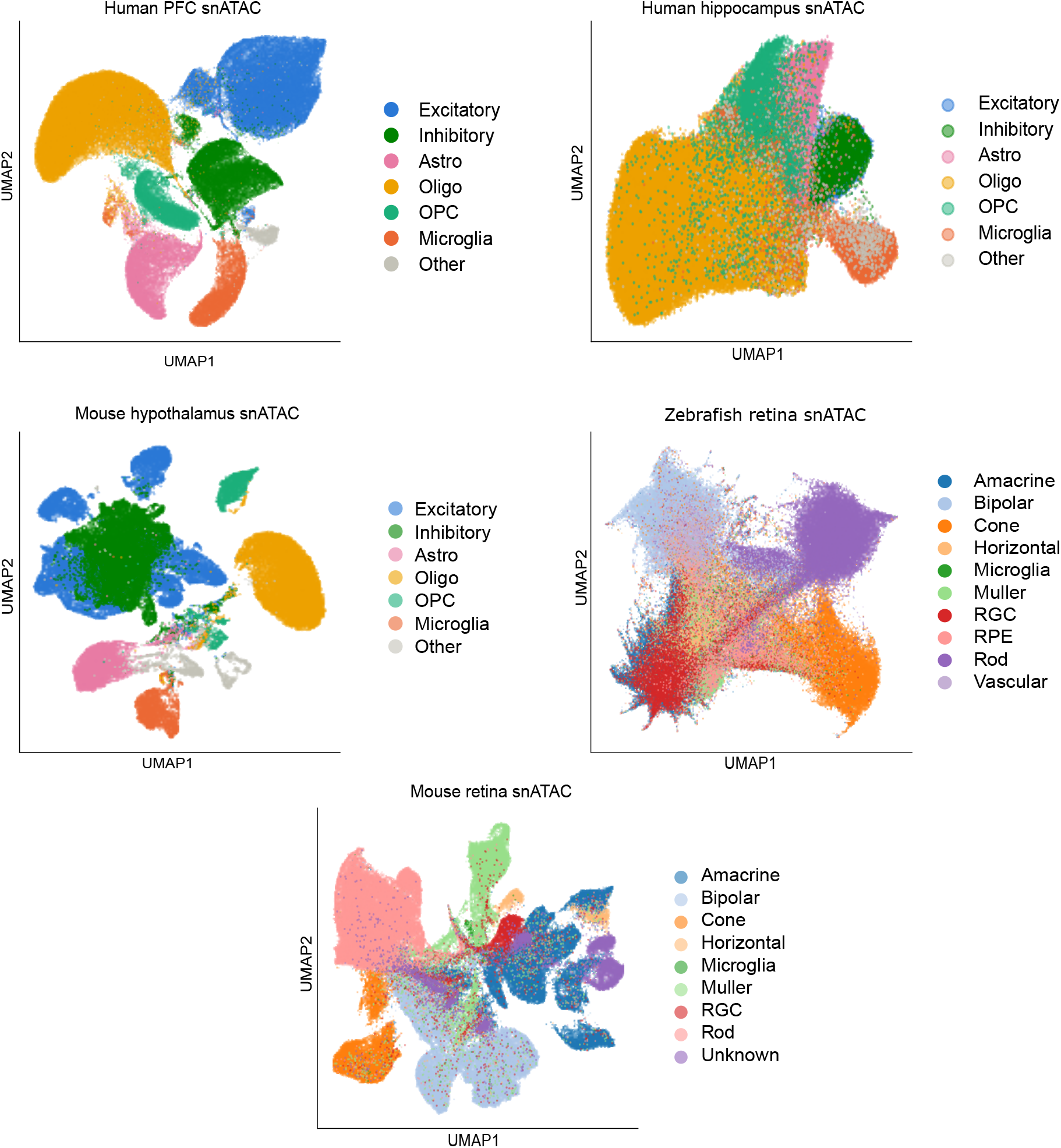
UMAPs for all healthy snATAC datasets, colored by the six major brain cell types or major retinal cell types.

**Supplementary Fig. S4:**
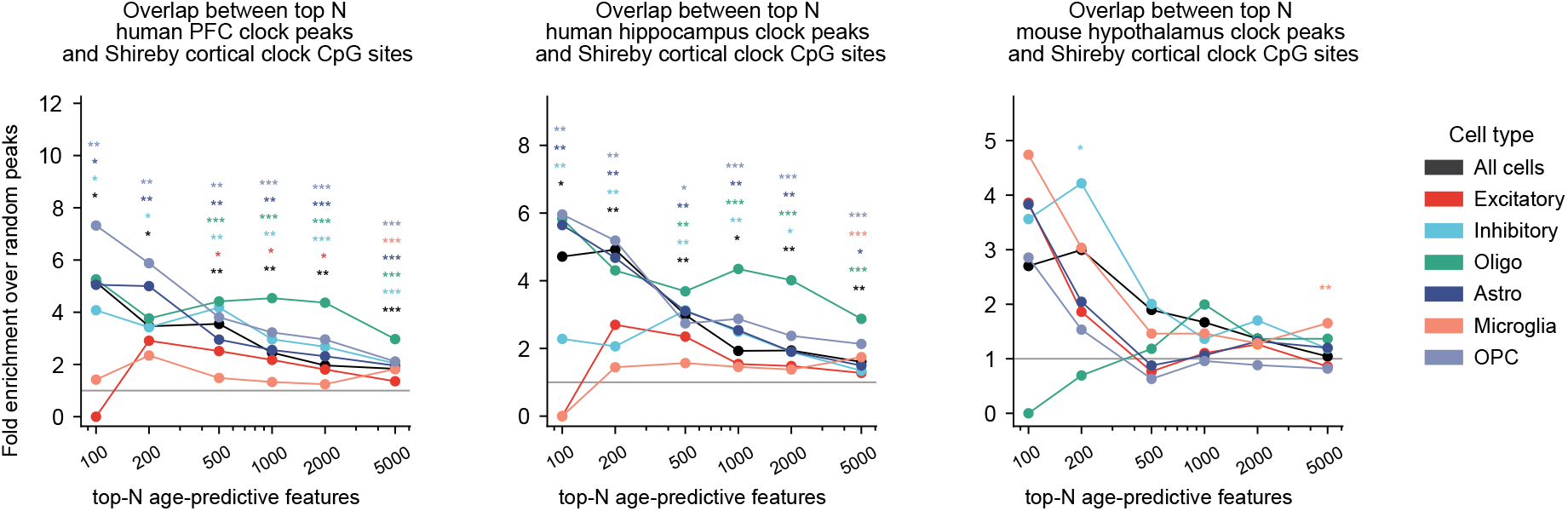
Enrichment of the top *n* PFC-predictive peaks and top *n* hippocampus-predictive peaks against 347 age-predictive CpG sites from a human cortical methylation clock, and enrichment of the top *n* mouse hypothalamus-predictive peaks against 435 age-predictive CpG sites from a mouse multi-tissue methylation clock.^91,92^ A snATAC-seq clock peak is considered to overlap a methylation clock CpG site if their midpoints are within 10 kb of each other. For each cohort, the top *n* age-predictive peaks were then tested by a hypergeometric test for overlap with age-predictive CpG sites, relative to randomly sampled peaks.

**Supplementary Fig. S5:**
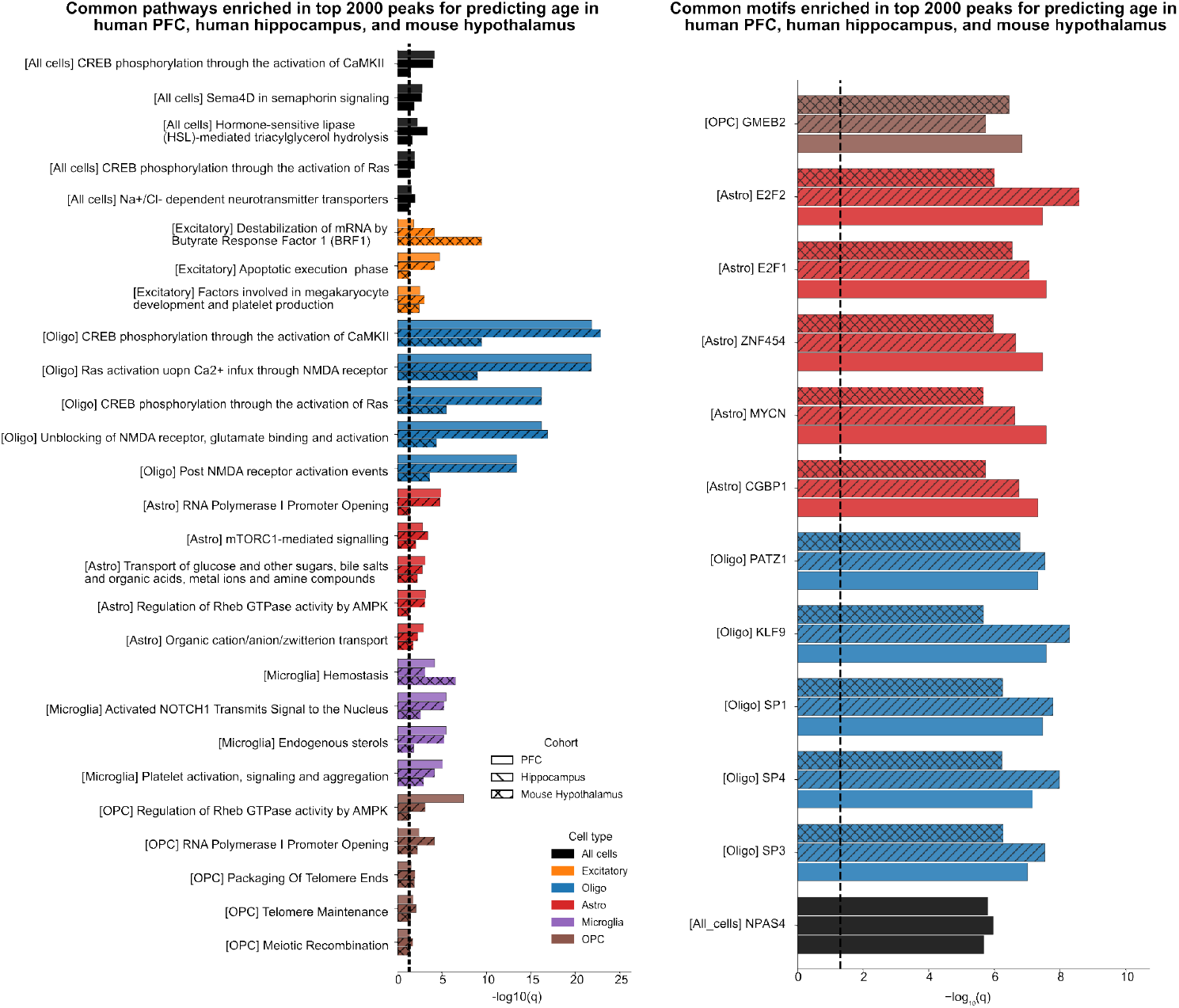
*Left*: Shared Reactome pathways enriched in the top 2,000 peaks across all three cohorts, by GREAT enrichment tests. *Right*: Shared HOCOMOCOv14 motifs enriched in the top 2,000 peaks across all cohorts, by an AME motif analysis.

**Supplementary Fig. S6:**
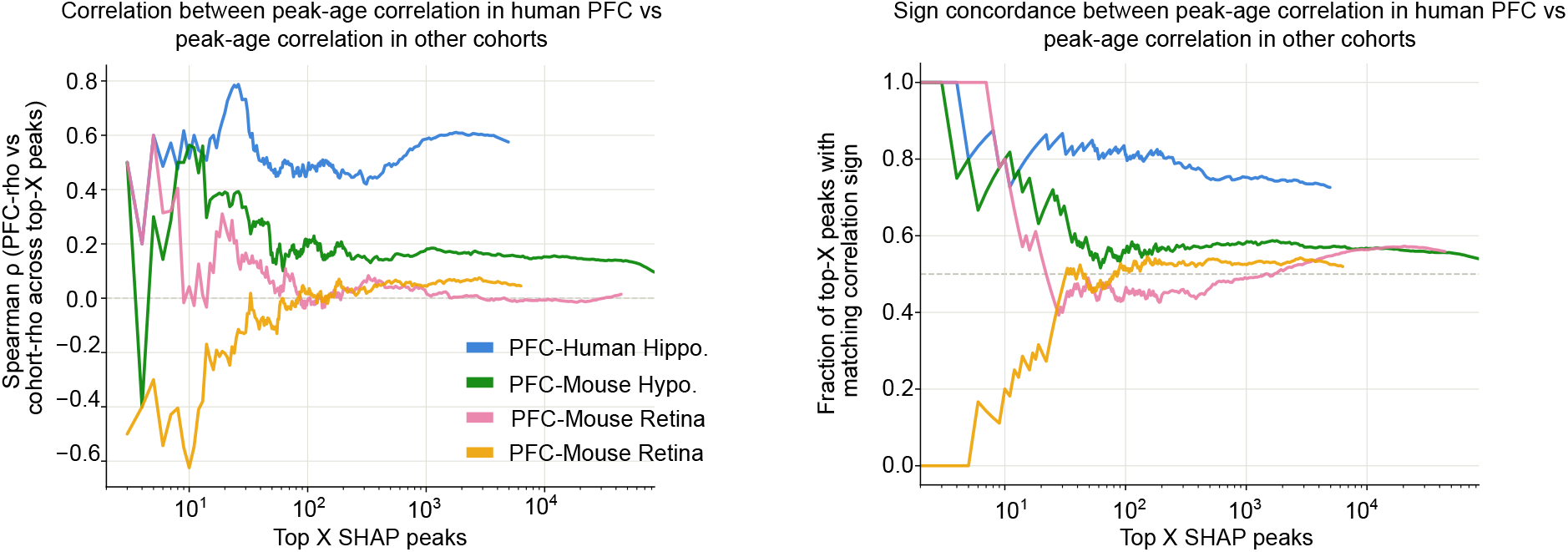
Relationship (Spearman correlation and sign concordance) between the peak-to-age correlation in healthy PFC and the peak-to-age correlation in other cohorts, across the top *n* PFC-predictive peaks that were also found in the other cohort.

**Supplementary Fig. S7:**
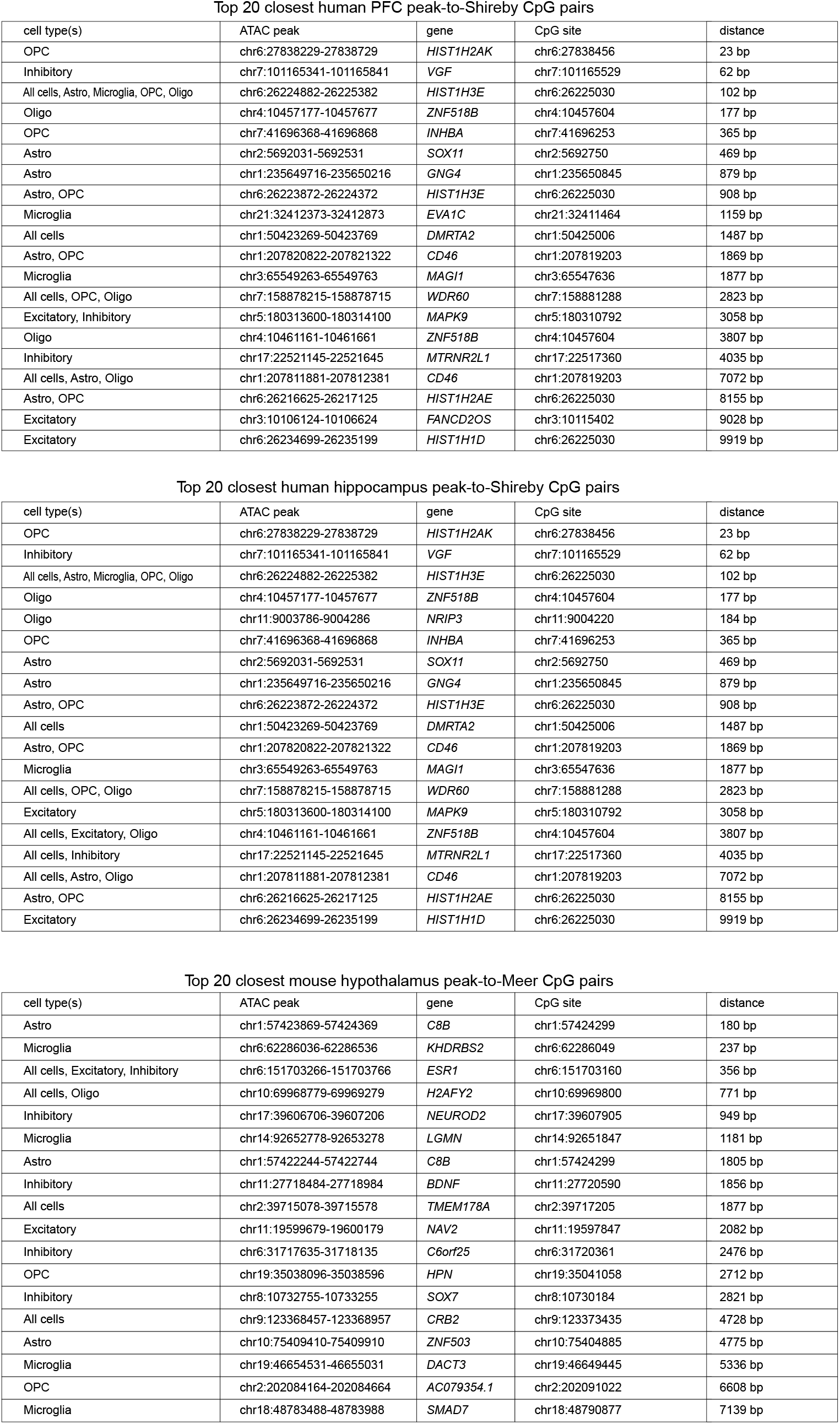
Top 20 peaks most proximal to methylation clock CpG loci, for each cohort.

**Supplementary Fig. S8:**
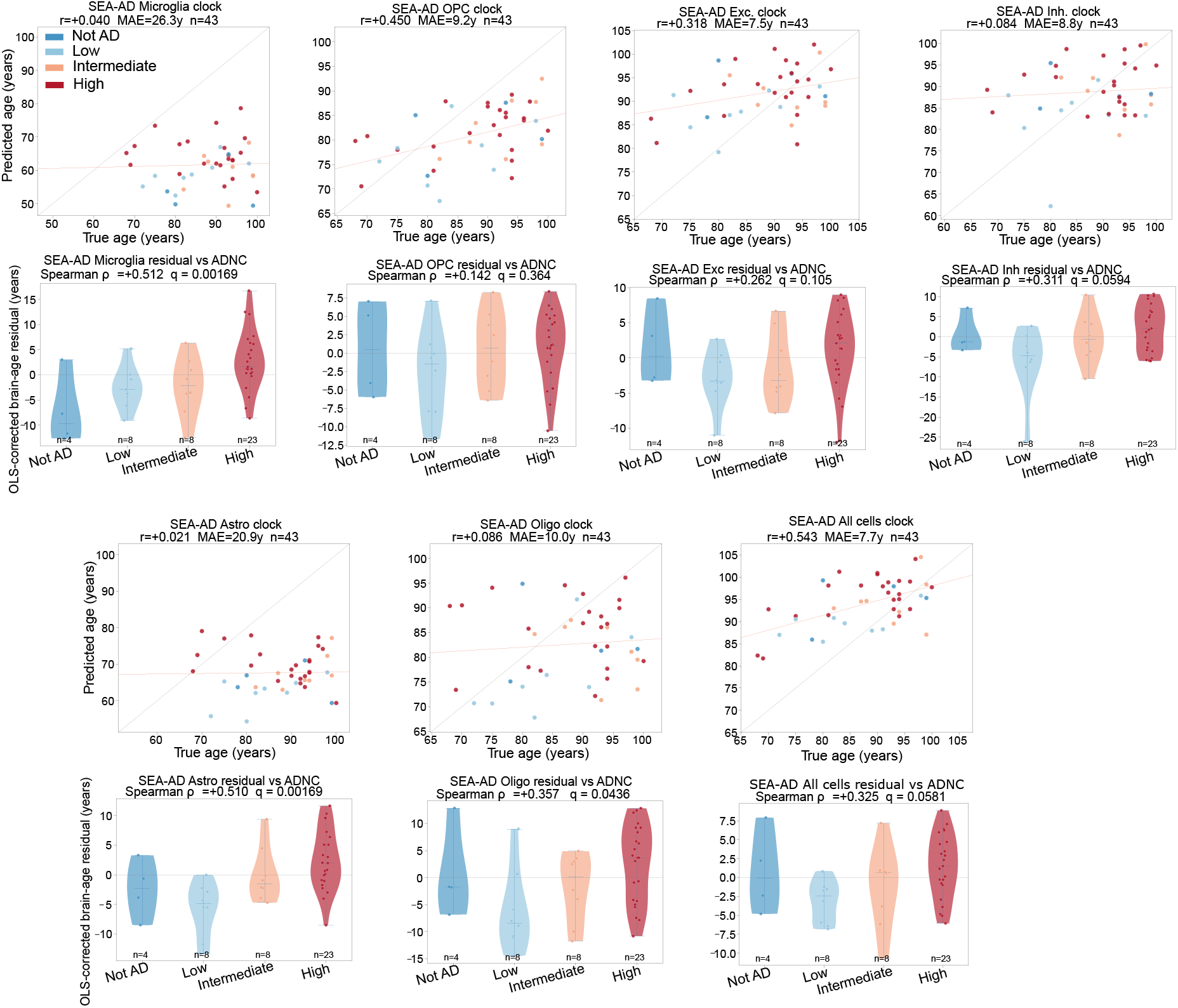
Healthy human PFC clock predictions on 43 SEA-AD DLPFC donors, for all cells and per cell type. Scatter plots show age predictions colored by ADNC status, and violin plots show age residuals for each ADNC category. The Spearman p-values are corrected for tests across multiple cell types.

## Notes

### Competing Interest Statement

The authors have declared no competing interest.

